# A Horizontal View of Primary Metabolomes in Vegetative Desiccation Tolerance

**DOI:** 10.1101/2023.02.10.528018

**Authors:** Halford J.W. Dace, Robbin Reus, Celeste Righi Ricco, Robert Hall, Jill M. Farrant, Henk W.M. Hilhorst

**Author notes:** Corresponding authors: Jill Farrant, Henk Hilhorst,.

## Abstract

Vegetative desiccation tolerance (VDT), the ability of such tissues to survive the near complete loss of cellular water, is a rare but polyphyletic phenotype. It is a complex multifactorial trait, typified by some near universal (core) factors but with many and varied adaptations due to plant architecture, biochemistry and biotic/abiotic dynamics of particular ecological niches. The ability to enter into a quiescent biophysically stable state is what ultimately determines desiccation tolerance. Thus, understanding of the metabolomic complement of plants with VDT gives insight into the nature of survival as well as evolutionary aspects of VDT. In this study we measured the soluble carbohydrate profiles and the polar, TMS-derivatisable metabolomes of 7 phylogenetically diverse species with VDT, in contrast with 3 desiccation sensitive (DS) species, under conditions of full hydration, severe water deficit stress, and desiccated.

Our study confirmed the existence of core mechanisms of VDT systems relying on either constitutively abundant trehalose, or the accumulation of raffinose family oligosaccharides and sucrose, with threshold ratios conditioned by other features of the metabolome. DS systems did not meet these ratios. Considerable chemical variations among VDT species suggest that similar stresses, e.g. photosynthetic stress, are dealt with using different chemical regimes. Furthermore, differences in timing of metabolic shifts suggest there is not a single “desiccation programme”, but that subprocesses are coordinated differently at different phases of drying. There is likely to be constraints on the composition of a viable dry state and how different adaptive strategies interact with the biophysical constraints of VDT.

## Introduction

Plant vegetative desiccation tolerance (VDT), observed in so-called resurrection plants, is a rare and phylogenetically scattered example of anhydrobiosis, or life without water (Gaff and Oliver 2013). Many plants produce desiccation tolerant (DT) reproductive structures, but leaves, stems and roots must mitigate a wider array of stresses under extreme water deficit than the relatively simple structures of pollen, spores and seeds (Farrant et al. 2020). Such stresses include the mechanical loss of turgor and associated tension on plasma membranes, the loss of structural water essential for membrane and protein integrity, rising ion concentrations and macromolecular crowding and, due to the presence of photosynthetic metabolism, massive photooxidative damage as water is removed (Bartels and Hussain 2011; Farrant et al. 2020). The associated adaptations have been much studied but questions remain about the exact parameters of the small molecule and macromolecular compositions that make anhydrobiosis possible.

The survival of near-total water loss and the ability to restore normal metabolism on rehydration require multiple strategies to mitigate such diverse stresses. Some of these stresses, and hence mitigations, are ubiquitous to DT systems; they would have been encountered by the first ancestral proto-plants to colonise land (Porembski 2011; Gaff and Oliver 2013). Near-universal strategies include ABA signalling (Cuming and Stevenson 2015), the accumulation of soluble sugars (Ghasempour et al. 1998), and the expression of Late Embryogenesis Abundant (LEA) family proteins (Artur et al. 2018). Other strategies depend on details of plant architecture, biochemical pathways and enzymes available to the species, and ecological niches with their particular biotic and abiotic dynamics (Farrant 2000). These phylogenetically local adaptations also include leaf folding, the accumulations of particular antioxidants or redox reserves (Farrant et al. 2015), and the disassembly, or modification and protection of the photosynthetic apparatus (Tuba and Lichtenthaler 2011; Christ et al. 2014; Zia et al. 2016; Radermacher 2019) during dehydration. The combination of universal with local adaptations may explain why vegetative desiccation tolerance is observed in so many plant clades, yet in a small minority of land plants. Independent evolution in multiple clades may have occurred by co-opting existing / ancestral functional networks in combination with new adaptations to a specific niche or plant architecture (Gaff and Oliver 2013). Alternatively, a ubiquitous ancestral trait may have been lost multiple times in plant evolution. This emergence from a mix of universal and local sub-traits, is why traditional functional genetic analyses in model species have not yet proven useful for this complex, multifactor trait.

Early work contrasting orthodox (DT) and recalcitrant / desiccation sensitive (DS) seeds revealed phase transition thermodynamics compatible with the formation of vitreous phases in the cytoplasm (Vertucci and Leopold 1987; Vertucci and Farrant 1995). Subsequently, studies have suggested that the Natural Deep Eutectic Solvents (NADES) could form at specified molar ratios of certain metabolites, and may function as anhydrous solvents in the dry state (Choi et al. 2011; Oliver et al. 2020). More recently, it has been proposed that some LEA proteins may be involved in complex liquid-liquid phase separations that protect other molecular species during anhydrobiosis. It is entirely possible that multiple phase systems may be involved, either in different biological systems, in different cellular compartments, or at different water contents or temperatures (Belott et al. 2020; Toit et al. 2020).

The mass accumulation of soluble carbohydrates is a common feature among almost all anhydrobiotic systems (Ghasempour et al. 1998; Crowe 2014). In animal, fungal and nonvascular plants, trehalose and, in some instances, glucose have been reported to be accumulated to high levels (Ghasempour et al. 1998; Tapia et al. 2015). In angiosperm resurrection plants, sucrose and raffinose family oligosaccharides appear necessary for desiccation tolerance (Egert et al. 2015) (Farrant et al. 2015), while glucose and fructose become less abundant on drying (Ghasempour et al. 1998). The synthesis of raffinose is associated with modulation of the myo-inositol and galactinol pools, and significant up-regulation of the associated enzymes has been observed in DT Xerophyta spp (ElSayed et al. 2014).

Large shifts have also been observed in both amino acid profiles and the organic acids associated with primary carbon metabolism (Oliver et al. 2011). While these effects could be artefacts of both large shifts in the proteome and the combination of carbon reallocation and metabolic shutdown respectively, it is quite possible that the particular amino acid profiles and organic acid complements are adaptive and necessary to surviving water loss. Interestingly, the amino acid profiles observed in the context of desiccation tolerance differ from those observed under drought-stress more broadly, with milder water deficit stress inducing the accumulation of proline, presumably as an osmoprotectant, while VDT plants have been reported to accumulate other amino acids, including GABA, asparagine and tryptophan (Yobi et al. 2012; Dinakar and Bartels 2013; Radermacher 2019; Gabier et al. 2021).

Various studies have reported metabolomic responses to desiccation in individual resurrection plant species, but it remains difficult to make direct comparisons among data sets since metabolomic studies remain more susceptible to variations in methodology than, for example, genomics. We believe that a broad view of comparative metabolomics among a wide variety of resurrection plants has the potential to address a number of outstanding questions concerning the role of small molecules in anhydrobiosis and the mechanisms by which they act, and thus assist us in taking larger steps forward.

In this study we measured the soluble carbohydrate profiles and the polar, TMS-derivatisable metabolomes of a phylogenetically diverse set of resurrection plants, in contrast with selected DS species, under conditions of full hydration, severe water deficit stress, and desiccation to equilibrium with air. The DT species studied include the spikemoss *Selaginella lepidophylla*, the dicotyledons *Myrothamnus flabellifolia* and *Craterostigma pumilum* and the monocotyledons *Eragrostis nindensis*, *Xerophyta elegans, X. schlechteri* and *X. humilis*. DS systems included for contrast were *Arabidopsis thaliana* and the cereal grass *Eragrostis tef*, as well as desiccation-sensitive older leaves of *E. nindensis*. The goal of the study was to fill gaps in our understanding of the relationship between primary metabolomes and desiccation tolerance. In particular, we sought to address the following questions: (i) Do metabolite profiles suggest a mono- or polyphyletic origin for vegetative desiccation tolerance? (ii) Does metabolite profile data support the hypothesis that there are distinct early- and late-drying responses in DT plants? (iii) What are the constraints on the resurrection plant metabolome in the dry state, and do these constraints suggest particular model of biophysical adaptation?

## Materials & methods

### Plant cultivation and sampling

Individuals of each of the target species were grown in the greenhouse of the University of Cape Town, Rondebosch, South Africa with regular watering, having been sourced from the collection of that institution. After the harvesting of fully-hydrated samples, watering was withheld and plants were allowed to dry *in situ* until they reached equilibrium with ambient air.

Leaf tissue was sampled at full hydration, after severe wilting, and on desiccation. Leaves were dissected into two, with half being used for Relative Water Content (RWC) determination and the remainder of each leaf sample being frozen in liquid nitrogen immediately upon harvesting and later lyophilised prior to chemical analysis. RWC was determined gravimetrically as previously described.(Barrs and Weatherley 1962).

### Metabolite extraction

For soluble sugar analysis, 20mg of lyophilised leaf tissue was disrupted using a ball mill and extracted for 15 minutes in 1mL 80% methanol containing 400mg/L melezitose at 76 . Samples including leaf debris were dried *in vacuo* and resuspended in 1mL milli-Q water. After centrifugation, supernatants were further diluted 10x in milli-Q water for HPLC analysis. Since this method extracts 20mg of dry tissue per 10mL of HPLC-injectable sample, quantitative values in mg/L were divided by 2 to re-express them in mg / g dry weight.

For general metabolite analysis, small polar molecules were extracted from 10mg of milled, lyophilised leaf tissue by chloroform/methanol/water phase separation as described by Weckwerth *et al*. (Weckwerth et al. 2004) Extracts were dried *in vacuo* and sealed in GC-MS vials under argon.

### Ion-exchange chromatography

Soluble sugars were quantitatively determined using a Dionex ICS-5000+ DC ion exchange chromatography system equipped with a 4×250mm CarboPAC PA-1 column as previously described (Vidigal et al. 2016).

### GC-MS analysis

A GC-MS metabolomic analysis was conducted on dried polar extracts using a Thermo TSQ GC-triple quadrupole instrument. Samples were derivatised inline one-by-one with MSTFA immediately prior to their injection. Volatilised samples were separated on a TG5-MS capillary column and electron impact spectra were acquired in single-quadrupole mode. An alkane-based retention index ladder was added to each sample during the derivatisation step.

### Data pre-processing

GC-MS data were preprocessed with the R/Bioconductor version 3.10 metaMS package, using the package-native TSQXLS settings for peak detection and alignment.

Outlier detection was conducted using the ribitol internal standard, with samples whose ribitol peaks were more than 2 standard deviations from the mean removed from the analysis. Remaining peaks were standardised to the ribitol signal.

### Peak annotation

Peak spectra were extracted using MZmine 2.53, and masses rounded to whole values using the round-resampling filter before text mode export.

The retention times of alkane ladders were derived from extracted ion chromatograms for m/z = 57, and fitted to a Kováts retention model (Kováts 1958) based on defined alkane retention indices.

Peak retention indices were interpolated from the model and used in combination with spectra (preprocessed using MZmine 2.53). Preprocessed spectra and calculated retention indices were used to search the Golm Metabolome Database (Hummel et al. 2010) for annotation.

### Statistical analysis

Statistical analysis of the annotated peak area table was conducted in R. Significance testing was conducted per species and compound, using ANOVA and Tukey’s HSD in the case of groups with three treatments, and 2-tailed Student’s t-test in the case of groups with two treatments.

### Pathway Inference

Differential pathway regulation was inferred based on the KEGG Pathway Database using the PAPi package (Aggio et al. 2010) from Bioconductor 3.10. Pathway predictions were filtered to exclude pathways not known in plants, with remaining differential regulation predictions serving as hypotheses for the interpretation of subsequent transcriptomic experiments.

## Results

### Ion exchange chromatography of carbohydrates

Key results of the targeted analysis of soluble sugars are illustrated in Figure 1, with fuller values tabulated in supplementary table S1. Quantitative analysis revealed common features in the stress response of all angiosperm species sampled.

**Figure 1.**
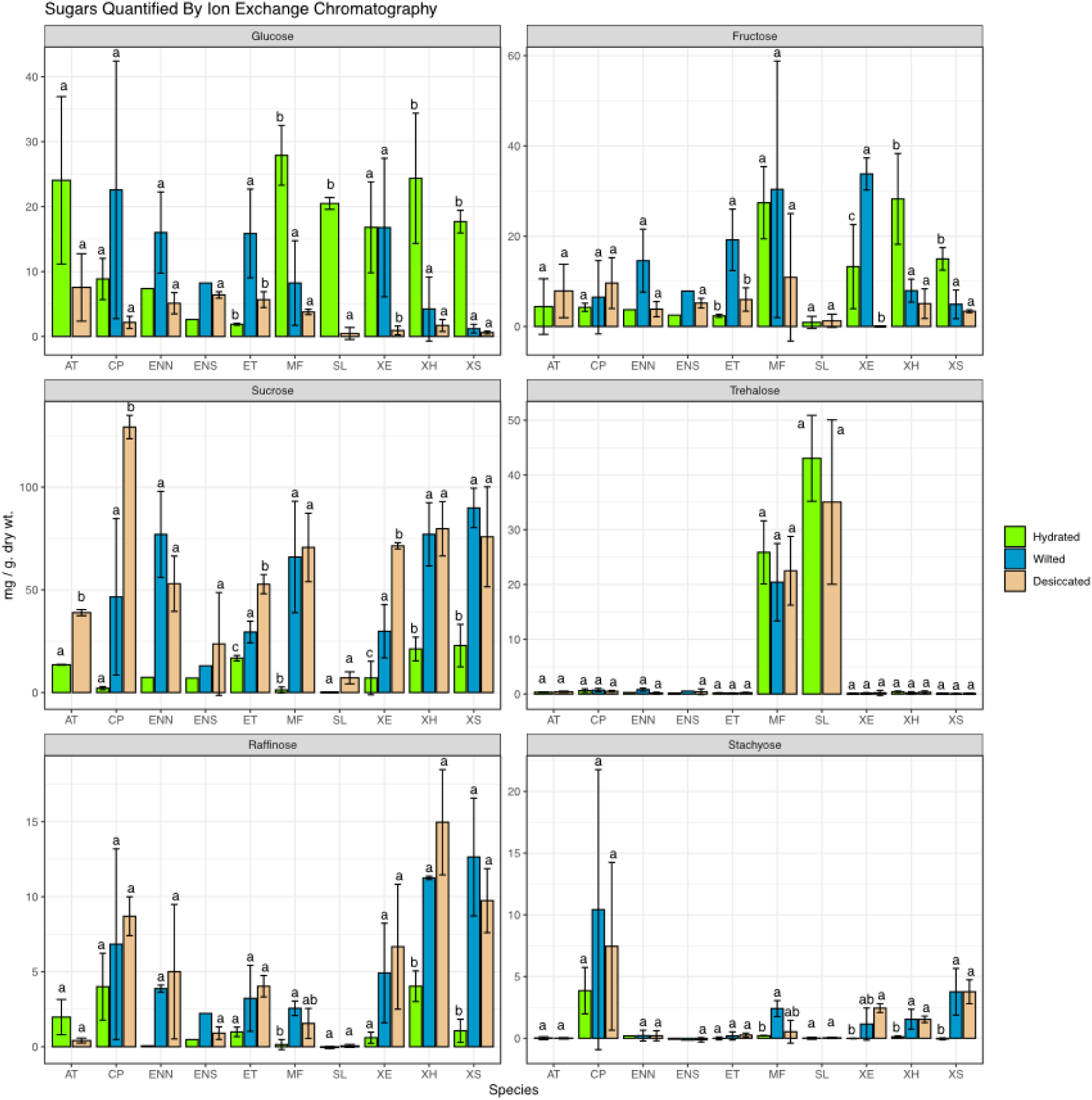
Abundance of soluble sugars measured by ion-exchange chromatography. Species: AT *Arabidopsis thaliana*, CP *Craterostigma pumilum*, ENN *Eragrostis nindensis* (Non-senescent), ENS *Eragrostis nindensis* (Senescent), ET *Eragrostis* tef, MF *Myrothamnus flabellifolia*, SL *Selaginella lepidophylla*, XE *Xerophyta elegans*, XH *Xerophyta humilis*, XS *Xerophyta schlechteri*. Significant differences determined by Tukey HSD. Bars with different letter annotations (a, b, c) within a single experimental group (sugar and species) have p < 0.05. n = 3 leaf samples from unique plants.

In all species, sucrose was accumulated during water deficit although the effect size was very small in *S. lepidophylla*, and in most (including all desiccation tolerant angiosperms), the monosaccharides glucose and fructose became less abundant over the same period.

The trisaccharide raffinose showed increases in abundance in the leaves of *C. pumilum*, all *Xerophyta* spp., the non-senescent leaves of *E. nindensis*, in *E. tef* and to a lesser extent in *M. flabellifolia*. Trehalose, by contrast, was present at constitutively high levels in both *M. flabellifolia* and *Selaginella*. It is notable that all DT tissues exhibited *either* large increases of raffinose abundance, *or* constitutively abundant trehalose. In the case of the desiccation sensitive species *A. thaliana*, the lack of raffinose accumulation is the only visible difference in this data set between it and desiccation tolerant species.. The other DS model, being the senescent leaves of *E. nindensis*, is characterised by extremely high biological variation, possibly due to the difficulty of correctly identifying leaves likely to senesce, and the variety of pathways they travel to organ death during the drying process.

Ratios of raffinose to sucrose, and trehalose to sucrose (w/w) were calculated for the air-dry state in each species (Table 1). In almost all cases, Raffinose Family Oligosaccharide (RFO)-dependent DT plants accumulated raffinose amounting to over 8% of the sucrose abundance, while trehalose-dependent DT plants accumulated trehalose at over 30% of the sucrose abundance. DS models reached neither threshold. The desiccation tolerant *C.pumilum* was an exception to these observations, accumulating slightly less raffinose in the dry state than the desiccation-sensitive but hardy grass *E.* tef. It should be noted that resurrection plants of the Linderneaceae, including *Craterostigma*, exhibit unusual carbohydrate metabolism and may be stabilised by unusual sugars such as octulose and sedoheptulose in the dry state (Zhang and Bartels 2017). Nevertheless given that it does increase its raffinose abundance on drying, it is likely to be employing a modified raffinose-dependent strategy, in which critical molar ratios for vitrification are affected by its octulose pool.

**Table 1.** Ratios of raffinose to sucrose, and trehalose to sucrose, expressed as a percentage, in the dry-state samples of each experimental species. Raffinose-dependent desiccation tolerant species are highlighted in green, trehalose-dependent species in yellow.

| Species | Raffinose : Sucrose ratio (dry) | Trehalose : Sucrose ratio (dry) |
| --- | --- | --- |
| *Arabidopsis thaliana* | 0.011 | 0.011 |
| *Craterostigma plantagineum* | 0.067 | 0.004 |
| *Eragrostis nindensis* (NS) | 0.094 | 0.004 |
| *Eragrostis nindensis* (S) | 0.038 | 0.018 |
| *Eragrostis tef* | 0.077 | 0.004 |
| *Myrothamnus flabellifolia* | 0.022 | 0.319 |
| *Selaginella* | 0.00 | 4.90 |
| *Xerophyta elegans* | 0.093 | 0.004 |
| *Xerophyta humilis* | 0.19 | 0.004 |
| *Xerophyta schlechteri* | 0.13 | 0.00 |

### GC-MS metabolomes

The metaMS workflow yielded a table of 373 peaks, whose contribution to the water loss response of each species is illustrated by Principal Component Analyses (PCA) in Figures 2 to 5.

**Figure 2.**
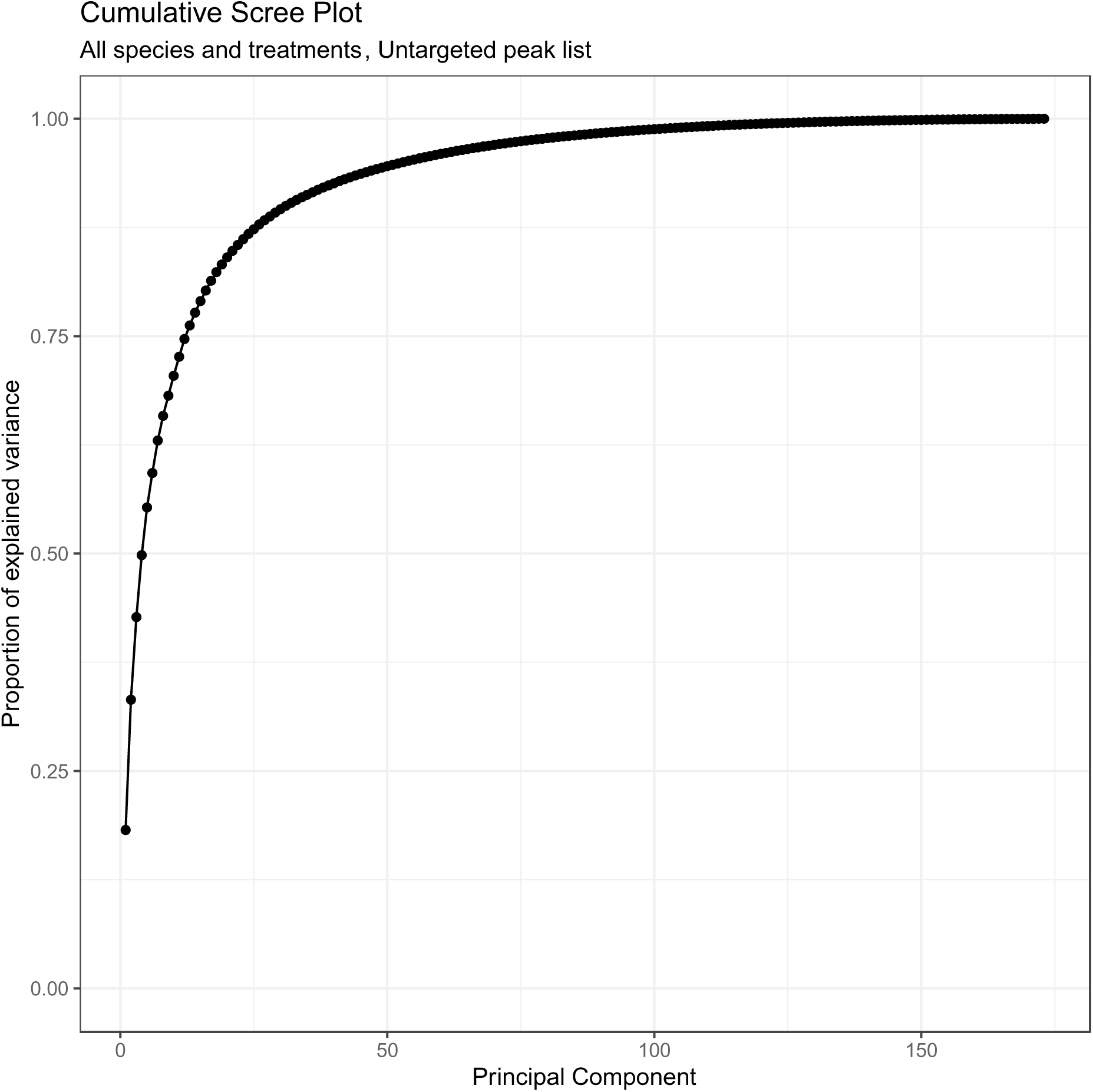
Cumulative explanation of variance, PCA conducted on all peaks returned by metaMS.

**Figure 3.**
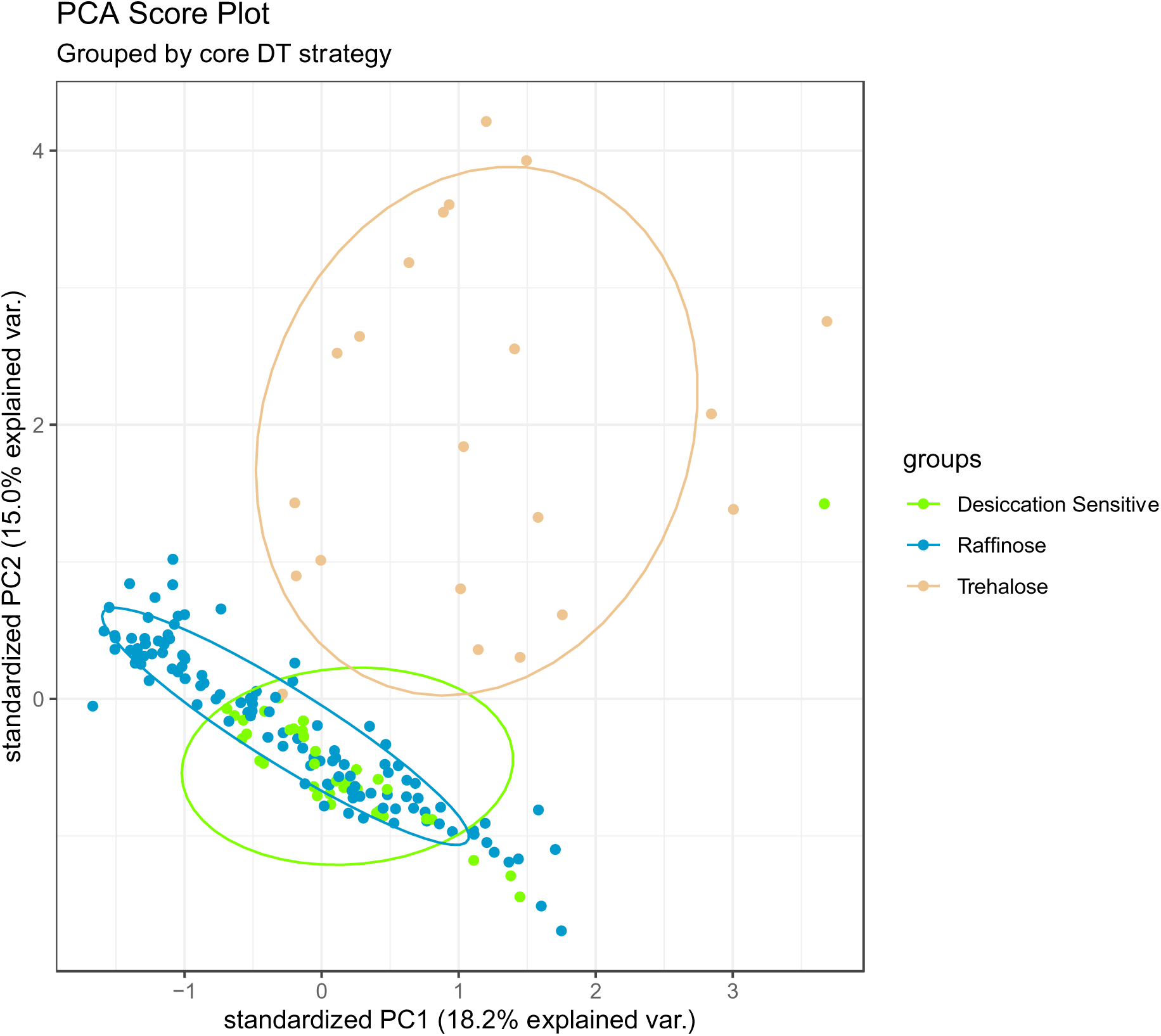
PCA score plot for all peaks returned by metaMS. Samples coloured by core desiccation tolerance mechanism.

**Figure 4.**
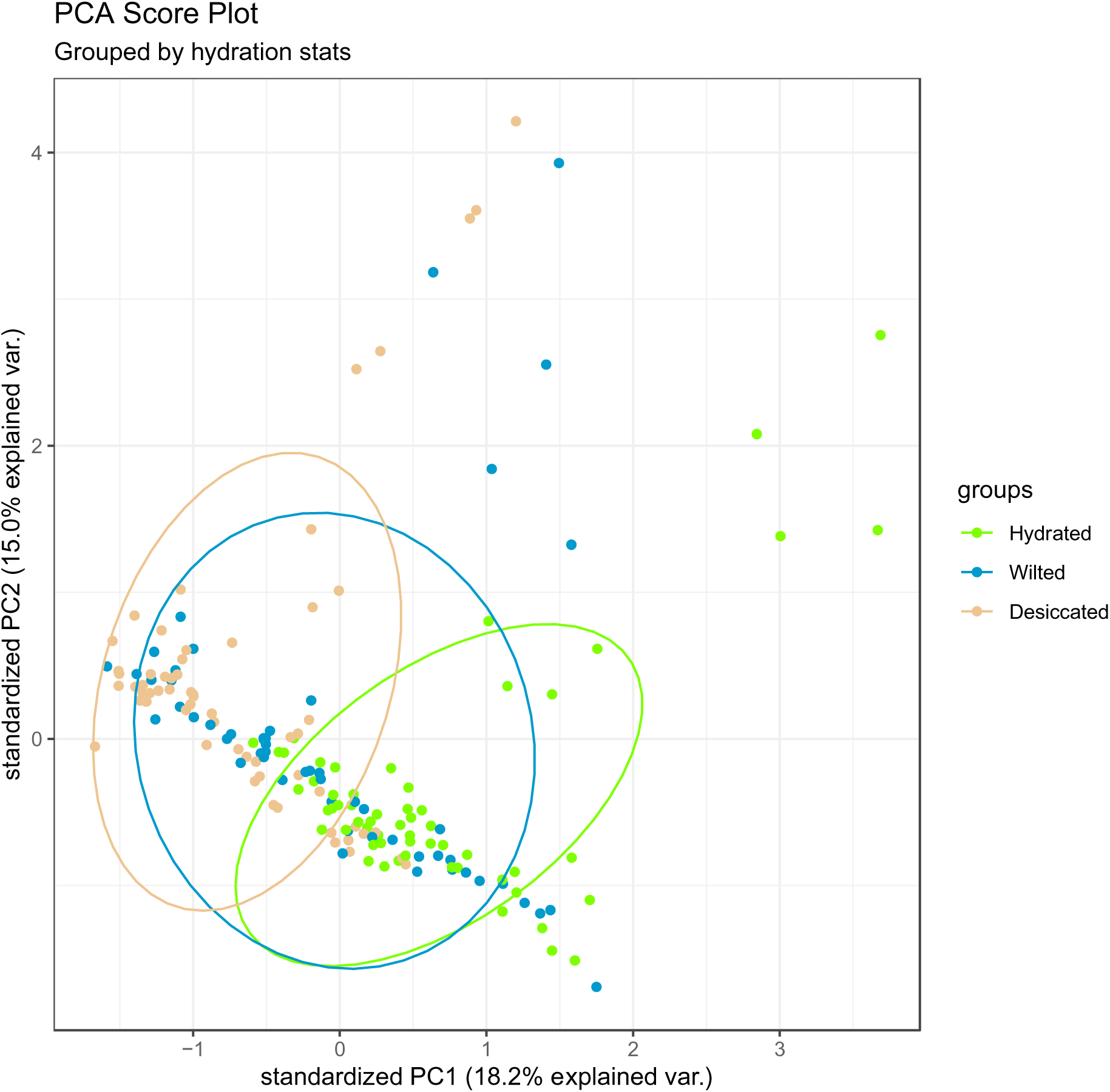
PCA score plot for all peaks returned by metaMS. Samples coloured by hydration status.

**Figure 5.**
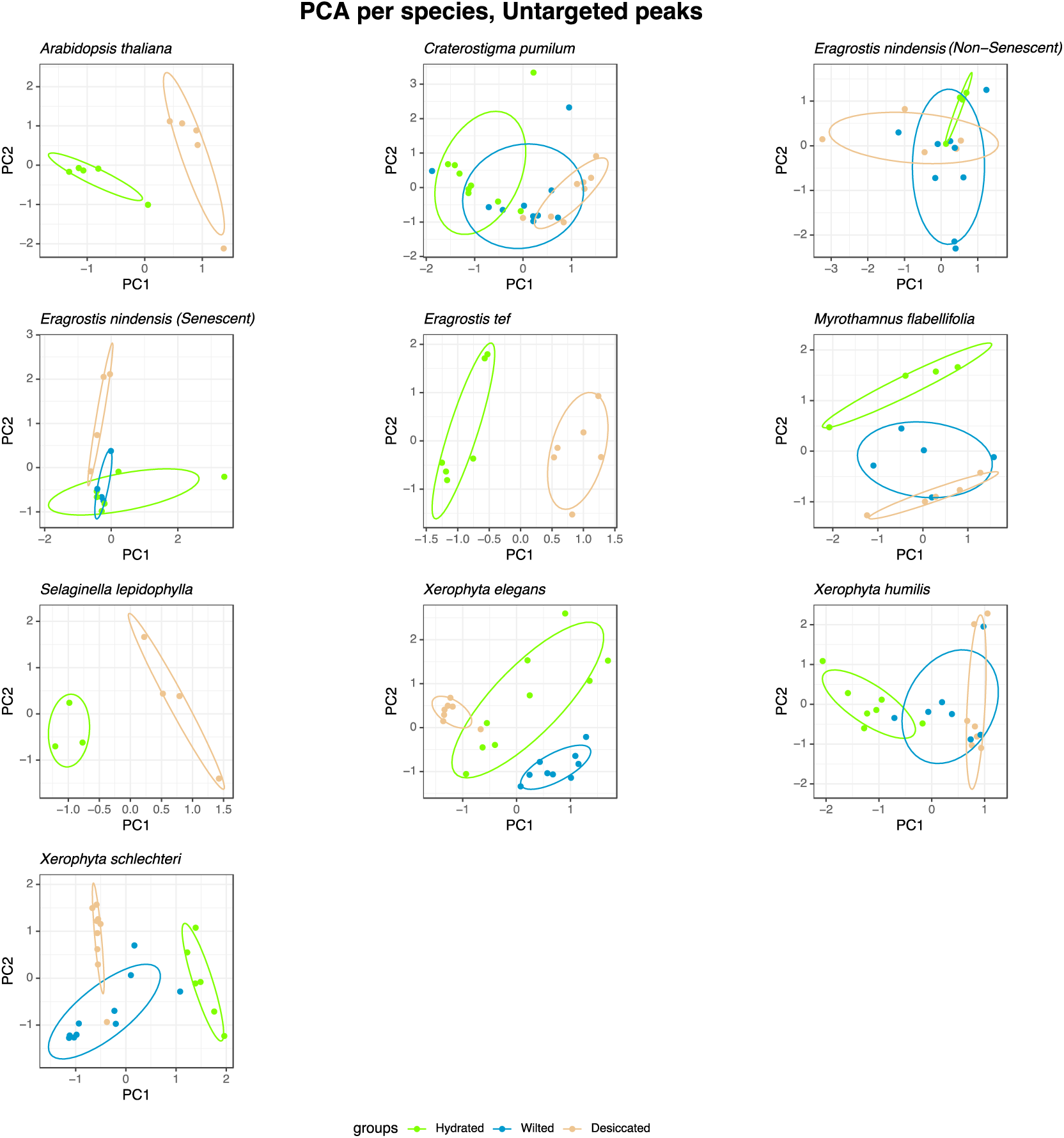
PCA score plot per species, all peaks returned by metaMS. Samples coloured by hydration status. Individual PCA calculations performed on mean-centered, Pareto-scaled data for each species.

Peaks most strongly associated with the drying response were annotated against the Golm Metabolite Database, producing a reduced subset of peaks, listed in Supplementary Table S1. Based on PCA analysis, the smaller peak set provided similar discriminatory power to the full feature set - see Supplemental Figures S1-S4.

Key selected observations:

The responses of various metabolite families to drying are illustrated in Figures 6-9.

**Figure 6.**
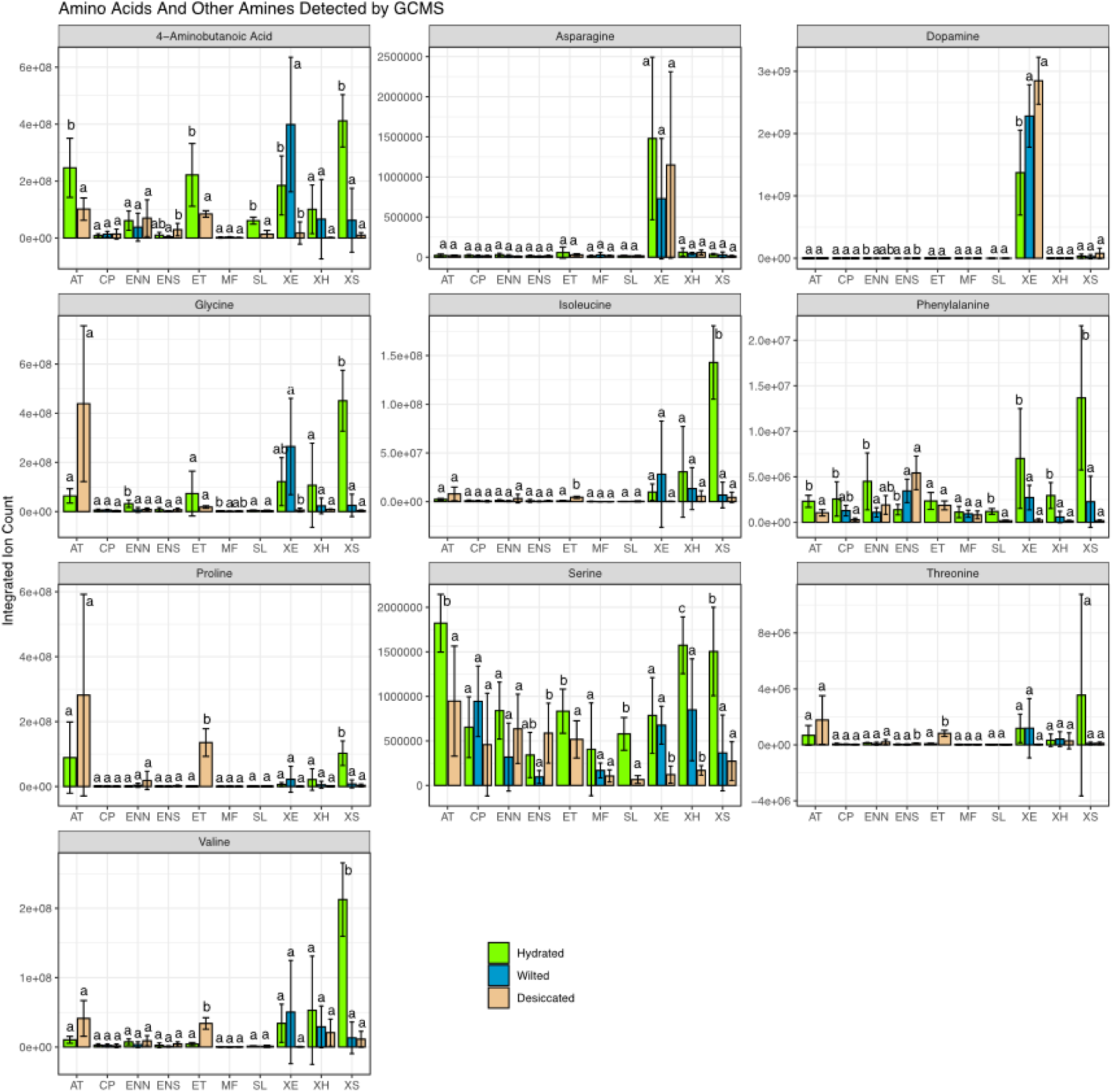
Desiccation response of amino acids and other amines detected by GCMS. Significant differences determined by Tukey HSD. Bars with different letter annotations (a, b, c) within a single experimental group (compound and species) have p < 0.05. n = 3 leaf samples from unique plants except where outlier removal reduced available sample pool.

Amino acids and other amines (Fig. 6) displayed diverse behaviours in the sampled species, with an overall tendency toward reduction in pools of free amino acids. It is noteworthy that proline, which is frequently cited as a compatible osmolyte involved in water deficit stress adaptation, accumulates only in DS *A. thaliana* and *E. tef.* Asparagine and dopamine (an amine included for its metabolic relationship with the aromatic amino acids), appeared almost exclusively in *X. elegans*, which otherwise has an unremarkable amino acid profile.

Metabolically core organic acids (Fig. 7) also showed diverse responses. The anomalous abundance of succinate in the hydrated samples of *C. pumilum* may be related to the unusual carbohydrate metabolism of that genus (Zhang and Bartels 2017).

**Figure 7.**
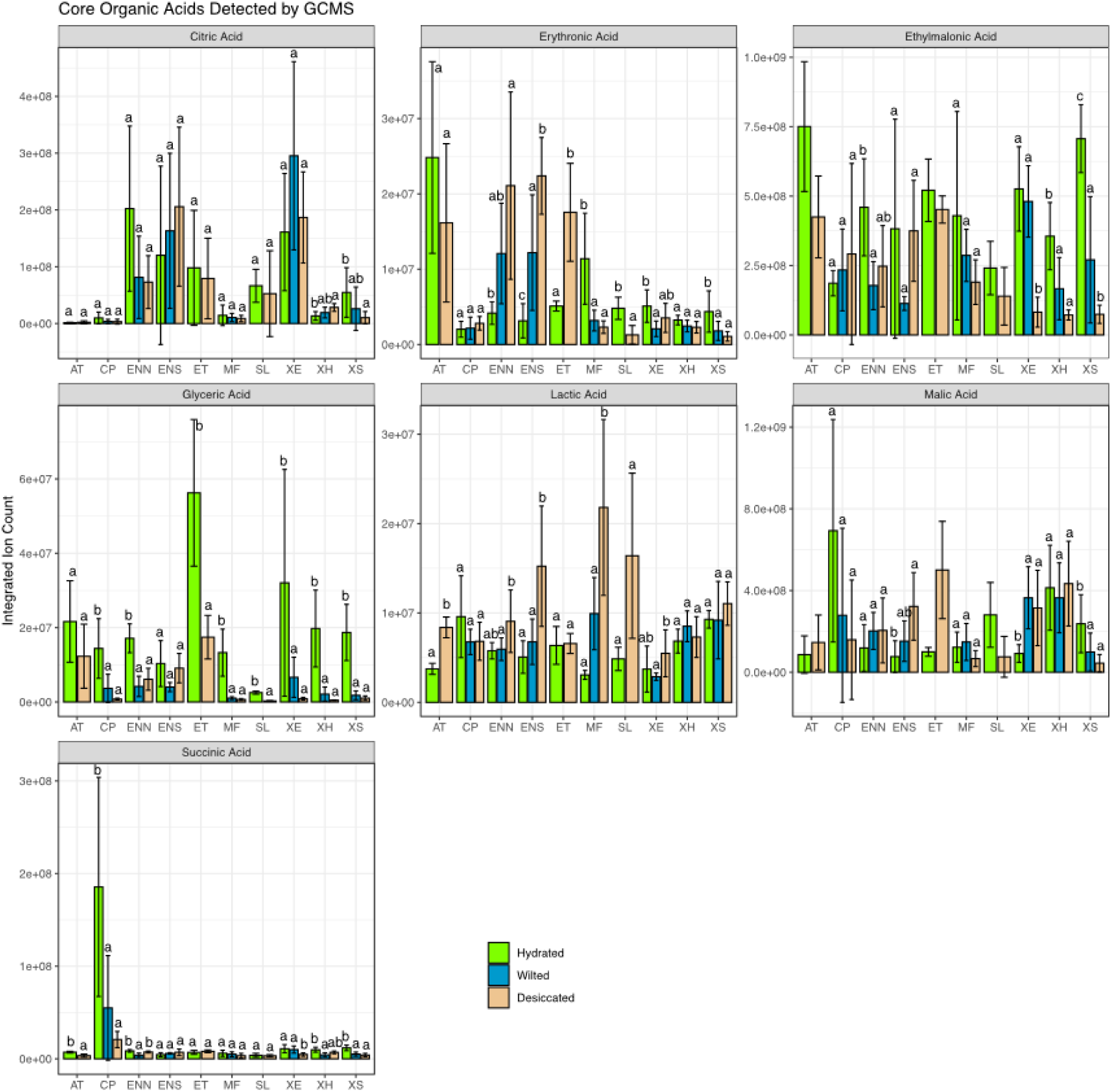
Desiccation response of organic acids detected by GCMS. Significant differences determined by Tukey HSD. Bars with different letter annotations (a, b, c) within a single experimental group (compound and species) have p < 0.05. n = 3 leaf samples from unique plants except where outlier removal reduced available sample pool.

It is notable that malic acid accumulated in the dry state most notably in the DS systems of *A. thaliana*, *E. tef* and the senescent leaves of *E. nindensis*; all DS systems. This accumulation may relate to the shutdown of core metabolism during water-deficit induced cell death.

Erythronic acid was observed to increase in abundance during drying in all *Eragrostis* systems, both DT and DS, but to show either little change, or decreased abundance in other systems. This suggests taxonomically specific lipid metabolism, rather than a specifically DT-related role.

Lactic acid showed little accumulation in most species, but more in the DS systems (*A. thaliana* and senescent *E. nindensis*), and considerably more in the robustly desiccation tolerant *M. flabellifolia* and *S. lepidophylla* — both trehalose-dependent resurrection plants.

Non-sugar compounds associated with carbon metabolism are highlighted in Figure 8.

**Figure 8.**
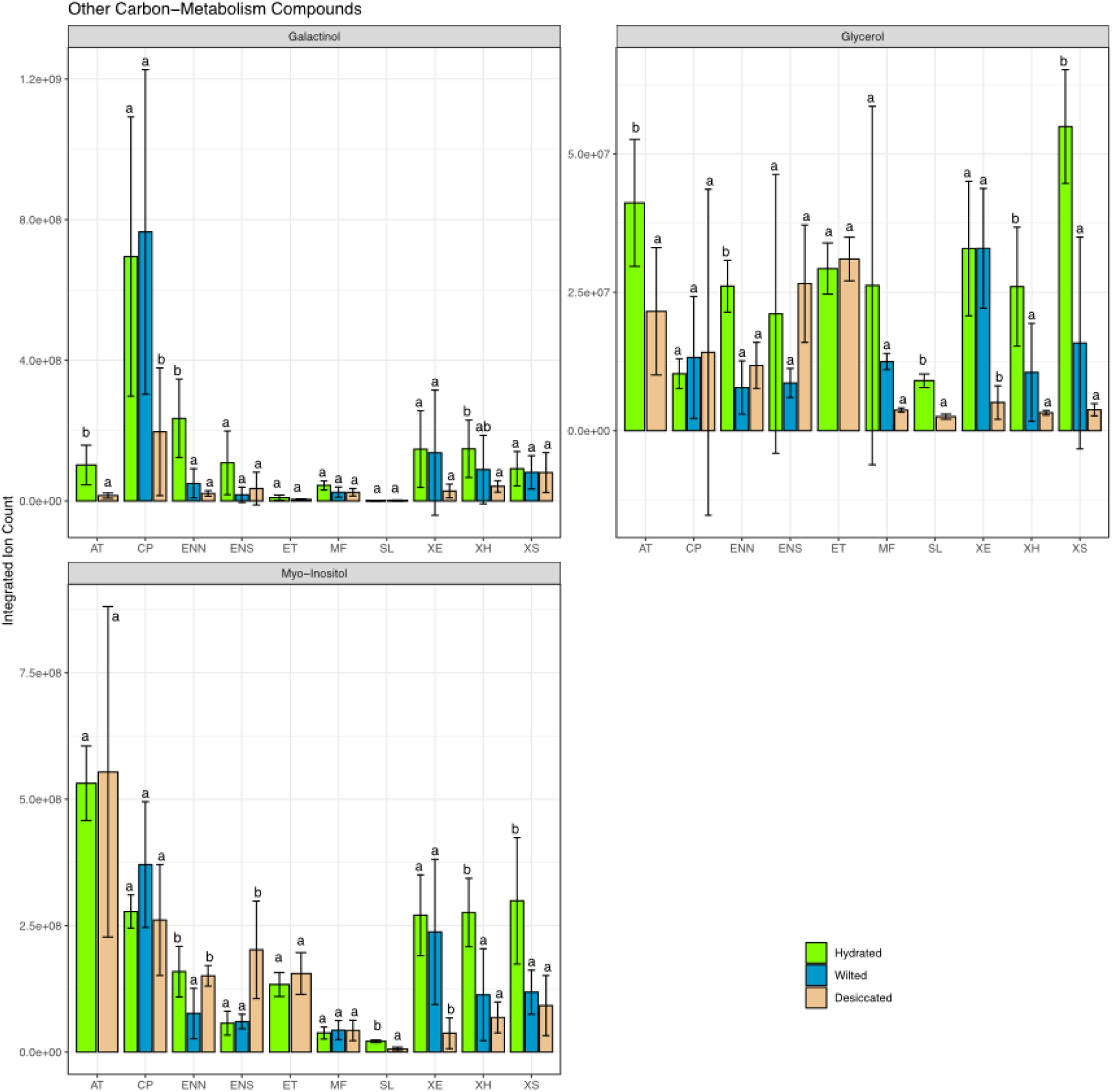
Desiccation response of carbon metabolites by GCMS. Significant differences determined by Tukey HSD. Bars with different letter annotations (a, b, c) within a single experimental group (compound and species) have p < 0.05. n = 3 leaf samples from unique plants except where outlier removal reduced available sample pool.

Most species that exhibited a strong raffinose response to drying also showed sharp reductions in galactinol pools, with implications for flux regulation of the raffinose synthesis pathway. Additionally, members of the *Xerophyta* genus showed sharp reductions in the abundance of both glycerol and myo-inositol, both involved in (*inter alia*) lipid metabolism, with this shift in abundance occurring earlier in *X. humilis* and *X. schlechteri* and later in *X. elegans*.

Compounds related to antioxidant metabolism are displayed in Figure 9. It should be noted that both gallic acid and quinic acid are observed at constitutively high levels in *M. flabellifolia*. This plant accumulates the polyphenol 3,4,5, tri-O-galloylquinic acid to very high levels in its vacuoles (Moore et al. 2005), and it is possible that the gallic and quinic acid signals may result from some hydrolysis of this compound during sample processing. Caffeoylquinic acid, an antioxidant chlorogenic acid, was present in all the *Xerophyta* species observed, although at higher levels in the poikilochlorophyllous species *X. humilis* and *X. schlechteri* and at lower levels in the homoiochlorophyllous, shade-dwelling *X. elegans*.

**Figure 9.**
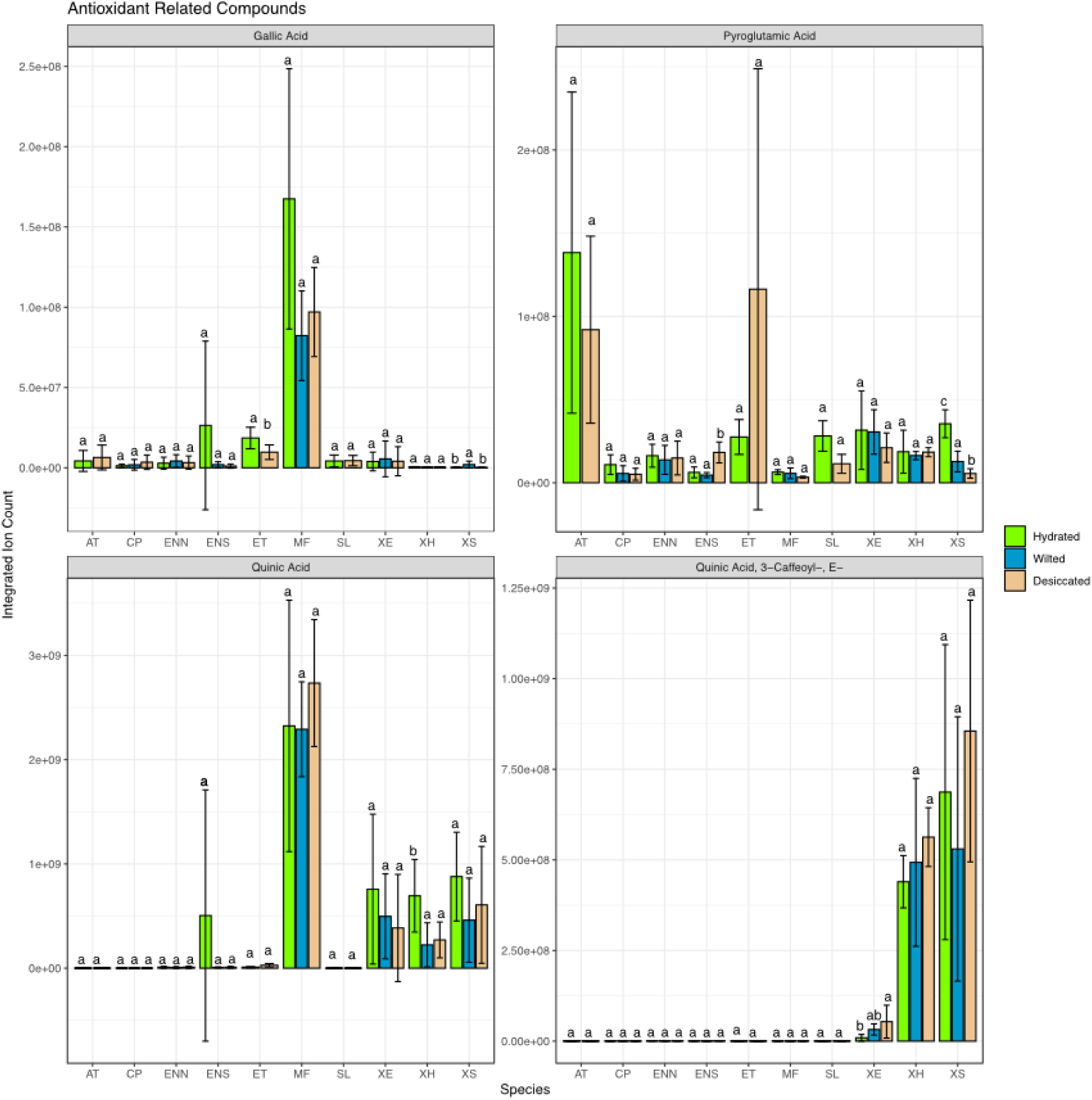
Desiccation response of antioxidants detected by GCMS. Significant differences determined by Tukey HSD. Bars with different letter annotations (a, b, c) within a single experimental group (compound and species) have p < 0.05. n = 3 leaf samples from unique plants except where outlier removal reduced available sample pool.

Total variance for the untargeted data set was measured (Fig. 10) in order to test the hypothesis that facultative (as opposed to constitutive) tolerance is associated with reduced variability in the dry state compared with the hydrated state. This hypothesis is compatible with the condition presumably imposed by the biophysical requirements for phase stability during anhydrobiosis. The pattern of reduced total variance in the dry state was evident in all DT species except *S. lepidophylla* (which is constitutively tolerant so presumably meets the conditions for anhydrobiosis at all times) and *E. nindensis*. It was also not observed in the desiccation-sensitive *A. thaliana*.

**Figure 10.**
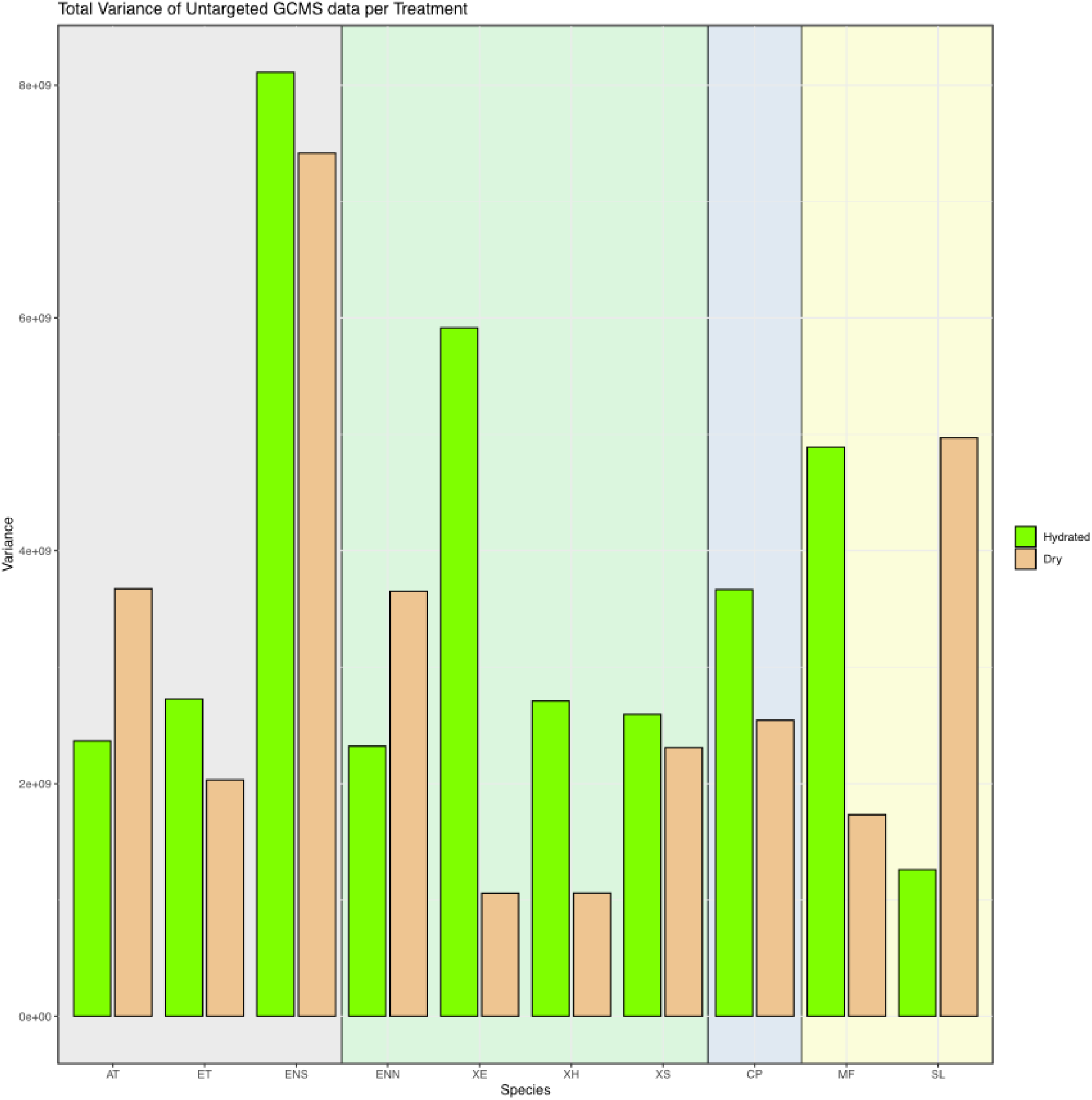
Variance per treatment and species. Total GCMS data set. Background colour illustrates core desiccation strategy: Grey – desiccation sensitive; Green – raffinose-dependent; Blue – raffinose modified by alternative carbohydrates; Yellow – trehalose-dependent. Total variances calculated from normalised, scaled, untargeted peak lists.

Hierarchical cluster analysis of the annotated data set (excluding sugars) revealed similarities in behaviour both in closely related groups of plants, and in closely related groups of compounds (Fig. 11). In the HCA output, related species clustered together, which all *Xerophyta* species exhibiting a similar pattern of metabolic shifts during drying, as did all members of the genus *Eragrostis*. Interestingly, given the omission of sugars from the HCA dataset, the two trehalose-dependent species *E. lepidophylla* and *M. flabellifolia* clustered together despite their very different phylogenetic contexts. DS models *E. tef*, *A. thaliana* and senescent *E. nindensis* do not cluster together, with *A. thaliana* forming a cluster with fellow dicots *C. pumilum* and *M. flabellifolia*, and spikemoss *S. lepidophylla*.

**Figure 11.**
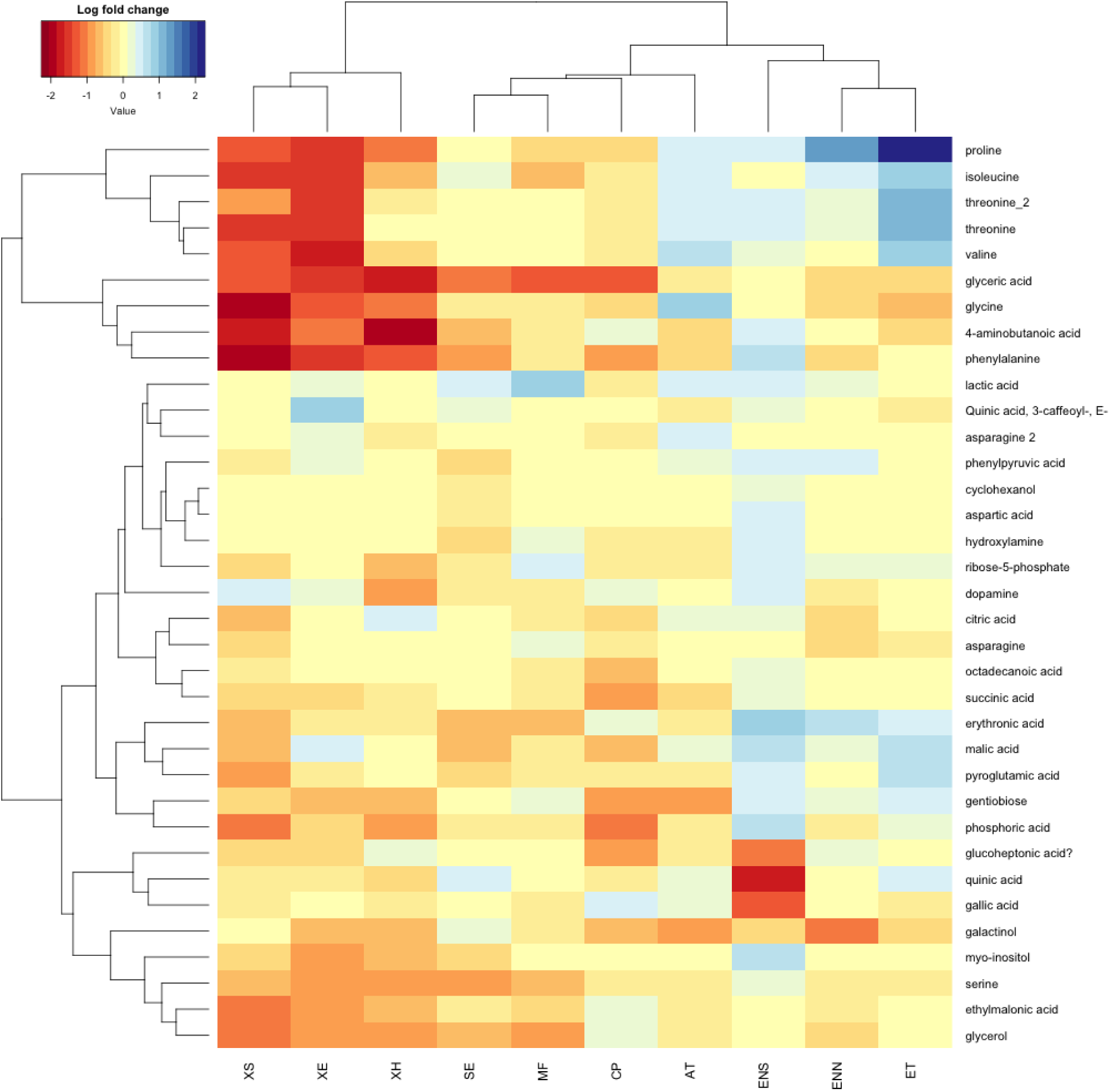
Cluster analysis based on change in abundance of targeted non-sugar compounds during desiccation. Heatmap colours represent Log_10_ change in mean abundance of each compound between fully hydrated and desiccated state. Cluster analyses on both axes calculated based on matrix Euclidean distance.

The clustering of compounds on the vertical axis reflected what appeared to be modules of common physiological regulation. For example all *Xerophyta* species exhibited decreased abundance in a cluster of amino acids including proline, phenylalanine, isoleucine, threonine and valine, while the *Eragrostis* grasses exhibit increases in abundance of the same cluster.

### Pathway Inference

The three pathways for each treatment most positively and negatively scored by the PAPi algorithm are listed in Table 2. Both positively and negatively scored pathways are selected as a high score magnitude, regardless of sign, representing a hypothesis that that pathway flux is modulated, with sign encoding information about the direction of proposed flux change.

**Table 2.**
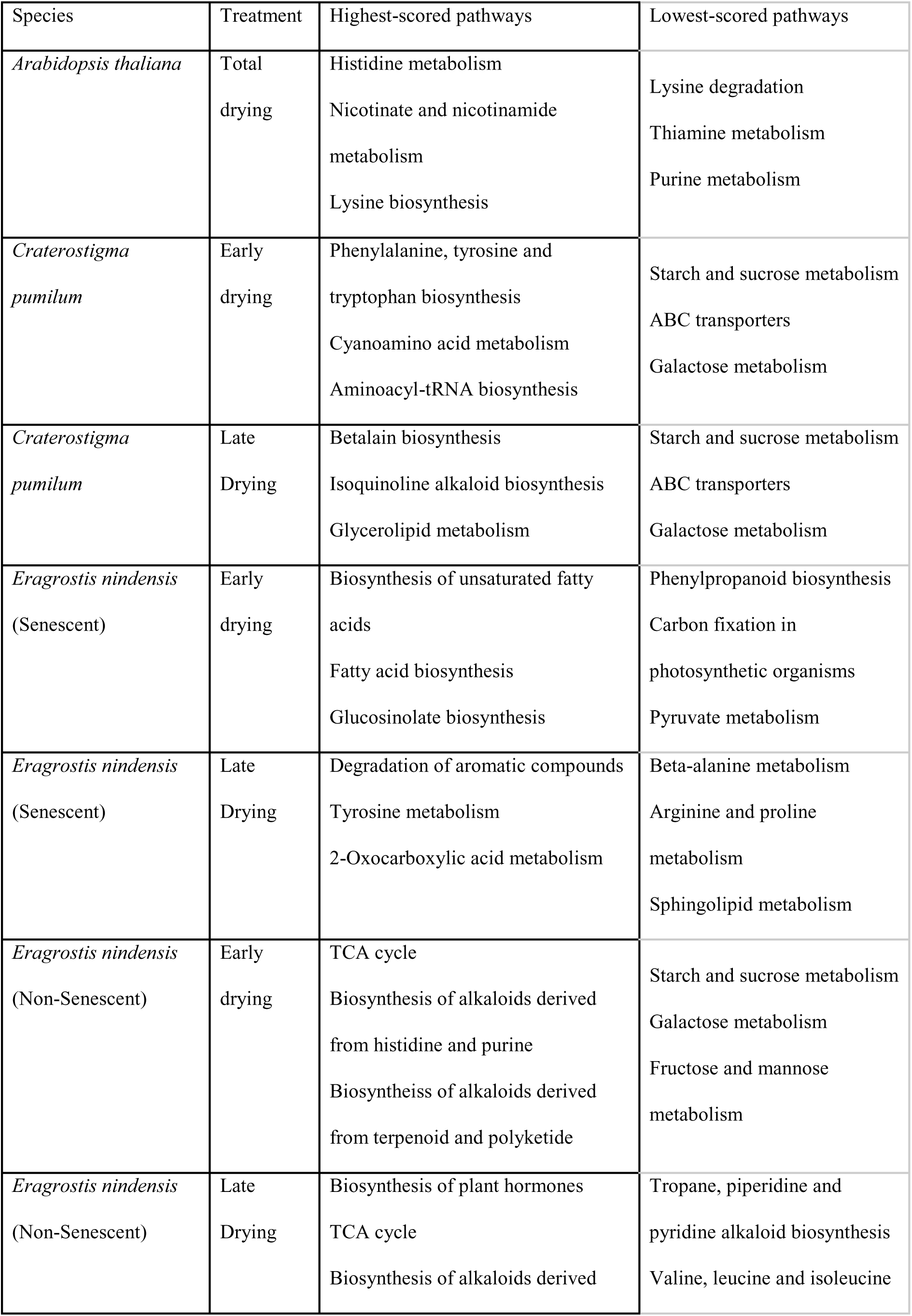
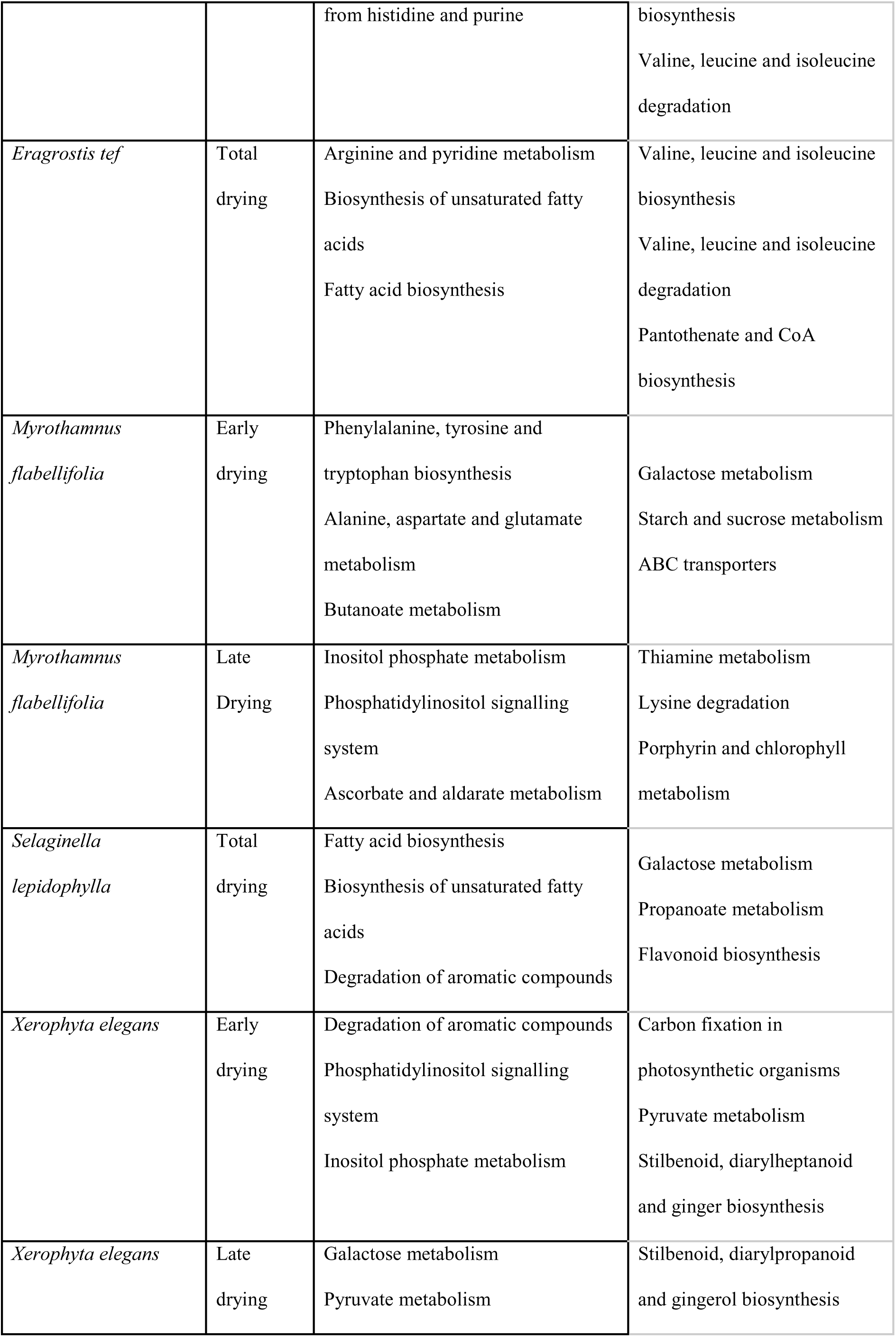
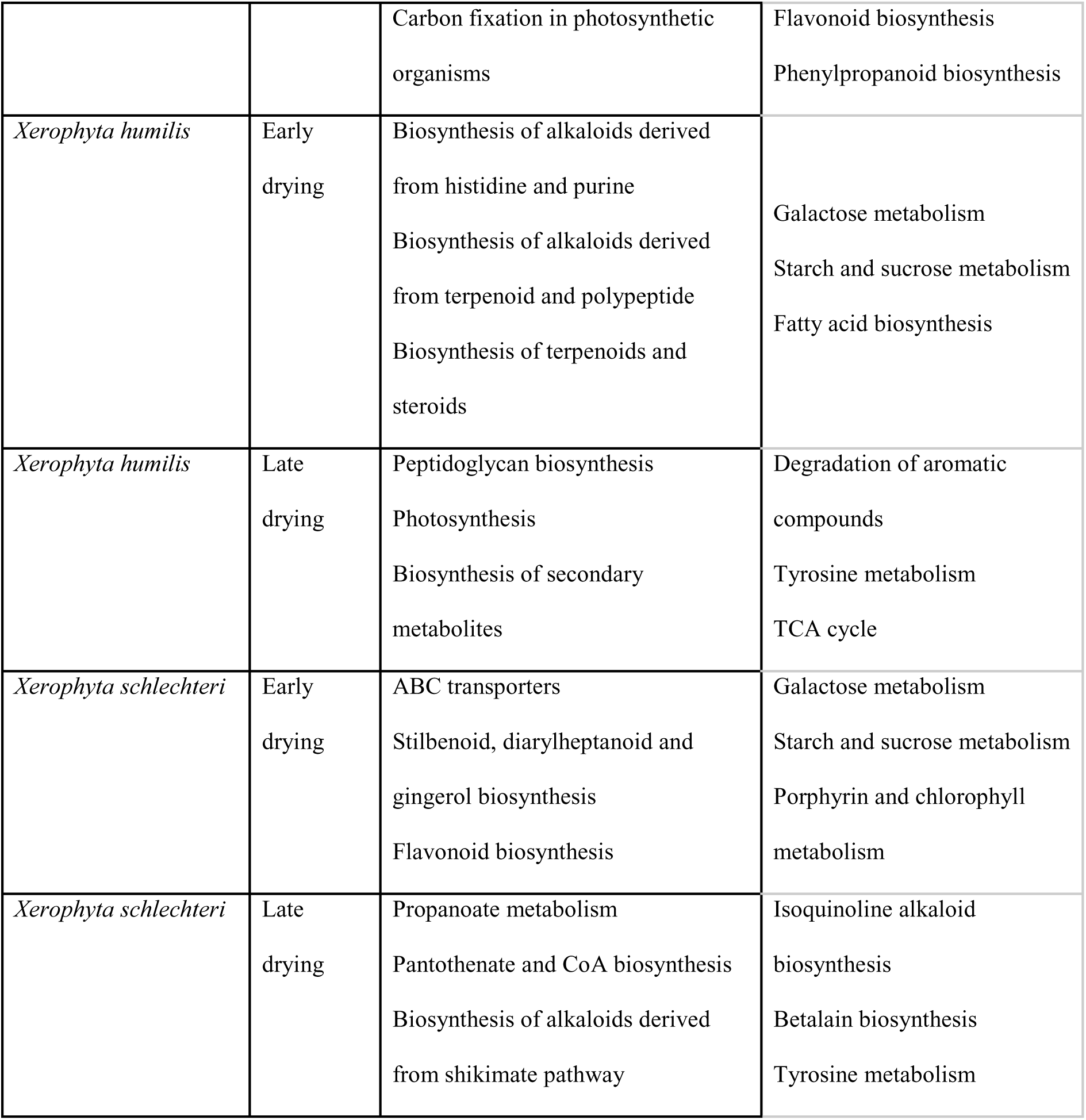
The three pathways for each treatment most positively and negatively scored by the PAPi algorithm are listed in.

Many species investigated yielded high-magnitude scores for carbohydrate pathways including Starch and Sucrose Metabolism, and Galactose metabolism, both key to the synthesis of key DT-related metabolites sucrose, raffinose and trehalose.

Numerous results pointed to a reconfiguration of lipids in drying systems, including pathways such as Fatty acid biosynthesis, Inositol phosphate metabolism and the phosphatidylinositol signalling system.

A contrast was observed between homoiochlorophyllous species such as *X. elegans* and its poikilochlorophyllous sister species *X. schlechteri*, with PAPi predicting significant regulation of porphoryn and chlorophyll metabolism (which includes chlorophyll catabolism) in the latter but not in the former.

Many predicted pathway activities point to modulation in the biosynthesis of secondary metabolites, which may include both antioxidants and antifeedants to protect the sugar-rich plants from herbivory in the dry state.

Desiccation-sensitive models *A. thaliana*, *E. tef* and senescent *E. nindensis* are enciched in pathways involving amino acid metabolism, which may be involved in attempts to conserve nitrogen from senescing tissues.

## Discussion

Analysis of the primary soluble carbohydrates and the TMS-derivatisable small molecule complement of a wide variety of resurrection plants revealed that all surveyed DT species depend either on constitutively high trehalose levels (in the case of more ancient lineages), or the adaptive accumulation of both sucrose and sufficient quantities of RFOs in more recent angiosperms. Additional critical functions in desiccation tolerance, such as maintaining redox equilibrium, appear to be fulfilled by a wider variety of mechanisms, and mediated by different metabolites, in different phylogenetic groups.

The existence of such core and non-core mechanisms poses several questions. As to the former, why the near-ubiquity of trehalose as a DT-enabler? And why have most angiosperm resurrection plants abandoned this otherwise universal mechanism for adaptive accumulation of sucrose and RFOS?

### Trehalose: Sufficient But Not Necessary?

Trehalose is a remarkable molecule, contributing adaptive utility to organisms across the tree of life. This disaccharide reduces glucose sensitivity in mice and humans (Yoshizane et al. 2020). It ameliorates physiological stresses in bacteria and confers desiccation tolerance upon yeasts (Tapia et al. 2015), brine shrimp, many tardigrades, nematodes and the midge *Polypedilum vanderplanki* (Okuda 2006), and even human cell cultures (Guo et al. 2000).

Among algae and non-angiosperm plants, trehalose is a vital part of both stress-related signalling and the bulk biophysical adaptation of cells to water loss (Avonce et al. 2006). In the latter role, it is hypothesised to act as a compatible solute and osmolyte under relatively mild water deficit, and as a water replacement medium and vitrification agent at very low water contents. Among angiosperms, trehalose has been reported to have a less significant role in adaptations to water loss (Gaff and Oliver 2013). In both orthodox seeds and those vegetative angiosperm tissues capable of anhydrobiosis, water replacement and vitrification seem to depend upon large accumulations of sucrose augmented by RFOs, primarily raffinose and stachyose. In the light of these prior observations, the current results raise some interesting issues concerning the role of these respective sugars in different DT systems, and the possible fitness advantage of either mechanism in different contexts.

### Induced versus Constitutive Adaptations

In all of those plants observed to use a trehalose-based stress response strategy, levels of trehalose are constitutively high, with relatively modest (if any) accumulations during the drying process. Even in the woody spermatophyte *M. flabellifolia*, which has ample time during drying to mount a metabolic response, we observed a strong, induced increase in sucrose abundance, but a very much smaller increase in the pool of already-abundant trehalose. By contrast, in those plants using a sucrose-RFO strategy, there appears to be a strongly induced character to the raffinose response. So we must ask, why is the sucrose-RFO strategy compatible with an induced response while the trehalose-dependent strategy appears relatively constitutive?

### The selective advantage of RFOs

With the exception of *M. flabellifolia*, all observed angiosperm resurrection plants eschew the otherwise-universal trehalose strategy in favour of the sucrose-raffinose strategy. This near-universality of the RFO strategy within such a large clade poses the question of why it has displaced the trehalose strategy so thoroughly (Sengupta et al. 2015). What is the selective advantage of galactosyl oligosaccharides over trehalose, given the latter’s effectiveness as an osmolyte, compatible solute and vitrifying agent?

The biosynthesis of raffinose by raffinose synthase (RafS) depends on the donation of a galactosyl moiety from galactinol, which is produced via the condensation of myo-inositol and UDP-galactose by galactinol synthase (GolS). Prior work on the evolution of this pathway shows it to be dependent on the evolution of GolS, which emerged early in evolution of higher plants (Sengupta et al. 2012).

RFOs have multiple roles in ameliorating abiotic stress in angiosperms (Obendorf 1997; ElSayed et al. 2014), however so has trehalose in other systems (Tapia et al. 2015). For example, both sugars have been identified as antioxidants. Nevertheless, the importance of antioxidant functions of individual species can be overstated given the general role of reduced carbon species in energy metabolism. It has not been established that they play a more significant antioxidant role than the major antioxidant systems of plant biochemistry, or that antioxidant potential is a major driver of selective pressure in the choice of carbohydrate regime.

In general, in order to increase fitness enough to precipitate a large-scale, stable evolutionary change across a large, diverse clade, a gene or regulon must either be more effective at achieving a certain physiological outcome (such as anhydrobiosis), or must be as good as its displaced predecessor while imposing lower costs on other aspects of fitness.

It would be difficult to argue that RFO-mediated anhydrobiosis is “more effective” than that mediated by trehalose. The ubiquity of trehalose-mediated strategies among species as diverse as baker’s yeast, tardigrades and *Selaginella* all point to the effectiveness of trehalose as an anhydrobiotic preservative. Equally, the extreme age reached by some angiosperm seeds in natural or artificial seed banks shows that raffinose-mediated DT can be more than sufficiently effective to meet the needs of plants growing and reproducing in natural environments.

It is plausible that the lactate accumulation observed in trehalose-mediated species in this study indicates an earlier shutdown of aerobic energy metabolism than that observed in the RFO/sucrose system. Nevertheless the robustness and longevity of the species in question in the dry state suggests that any such difference is not at a cost of survivability.

If RFOs are not “better” than trehalose, could they be metabolically “cheaper”? It is instructive to look at the relationship of sucrose, raffinose and trehalose abundances before and after drying in the set of sampled species.

All species exhibited increases in sucrose abundance during drying, to levels that ranged well within a single order of magnitude. No difference in sucrose accumulation is observed between DS and DT species, implying that sucrose abundance is not a sufficient explanation for the desiccation tolerance trait. Indeed, sucrose accumulation is observed in almost all plants in response to relatively mild water deficit, with sucrose acting as a non-toxic osmolyte. Thus while it forms a part of all plants’ drought response, this is distinct from the special requirements of anhydrobiosis.

With respect to the trehalose / raffinose dichotomy, it is worth noting that the lowest levels of sucrose in the unstressed condition were observed in *Selaginella* and *M. flabellifolia —* the two species which display constitutively high levels of trehalose and appear to rely on it for anhydrous glass formation.

Combined with this observation, the response patterns of trehalose and raffinose are quite different in species depending on these respective strategies: While raffinose increases during drying to levels that are modest in comparison with sucrose, trehalose is present at constitutively high levels in the two trehalose-dependent species, which *also* up-regulate sucrose abundance during early drying. This suggests that there is a requirement for constitutively high trehalose levels in trehalose-dependent organisms, by contrast with a relatively small requirement for induced RFO production in raffinose-dependent plants. The maintenance of abundant trehalose *in addition to* similar sucrose levels may effectively constrain the availability of carbon and reduction potential that could otherwise be deployed to growth or reproduction. Thus by evolving GolS and RafS, angiosperms may have given themselves a much more efficient route to balance the needs of growth and reproduction with the biophysical constraints of desiccation tolerance.

### Myrothamnus: An Exception To The Rule

Algae, early land plants, and extant nonvascular plants lack the sophisticated water-relations physiology of vascular plants, and spermatophytes in particular. Without deep root networks, and complex vascular and leaf tissues, such plants are subject to rapid hydrological equilibration with their environments, potentially drying and rehydrating at great speed. It makes sense that such plants should adopt a strategy of constitutive desiccation tolerance, since they may not have time to mount a complex, genetically programmed adaptive response during a drying event (Gaff and Oliver 2013). This implies that even if such plants possessed the biochemical elements of the RFO pathway, its relative utility in the context of desiccation tolerance would be marginal to negative, and constitutive trehalose remains as useful for them as it has for all other living kingdoms over millions of years.

The situation with angiosperm resurrection plants is quite different. Possessed of roots, complex xylem and refined management of leaf water efflux, they can retard water loss almost indefinitely, and have time for complex, induced responses to changing environmental and internal water potentials. Why, then, does *M. flabellifolia*, from the only known genus of woody resurrection plants (Sherwin et al. 1998), maintain constitutively high levels of trehalose (Drennan et al. 1993)? Although it is an ancient lineage, it is firmly within the angiosperm fold. This study reveals it to produce raffinose during drying, implying that it has a functional RFO pathway. Although galactinol was not detected in *M. flabellifolia* above the limit of detection by ion exchange chromatography, it was detected by the more sensitive GC-MS experiment. Although this needs to be confirmed via a genomic study, it is likely that *M. flabellifolia* possesses all the pathways necessary to pursue a raffinose-dependent DT strategy, yet the evolutionary fitness benefit appears to be much smaller than in other angiosperms. In addition to the genomic confirmation of how this species responds to water deficit stress, more ecophysiological research should be conducted to understand how its environment, unique growth habit and status as the only woody resurrection plant affect the dynamics of carbon and energy allocation to growth vs stress response (Marks et al. 2021).

### Craterostigma: A Distinctive Modification

*Craterostigma,* and other plants of the Linderneae, employ a distinctive and unusual carbon biochemistry, with members of this family storing large amounts of surplus sugar in the rare monosaccharide octulose, which appears to be used as a bulk carbon store in a way analogous to starch in other plants (Zhang and Bartels 2017). This sugar is likely to contribute to the anhydrobiotic stability of DT plants from this family, and to affect the remaining constraints on the molar ratios of remaining metabolites required for stable vitrification or NADES formation in *Craterostigma* spp. Further investigation into the physical chemistry of octulose may reveal why *C. pumilum* is able to survive the dry state with less RFO accumulation than other non-trehalose-dependent species.

### Xerophyta: Similarity With Difference

The three *Xerophyta* species surveyed in this study were substantially similar in their response to desiccation, and clustered together distinctly in the hierarchical cluster analysis. Yet they represent two quite distinct physiological patterns within VDT. *X. humilis* and *X. schlechteri*, which inhabit crevices on boulders, mountainsides and inselbergs and live in full sun, are poikilochlorophyllous, while the shade-dwelling *X. elegans* practices homoiochlorophylly (Tuba and Lichtenthaler 2011). Alongside this important difference, *X. elegans* appears to sequence its desiccation response quite differently from *X. humilis* and *X. schlechteri,* with sucrose accumulation, amino acid depletion and large shifts in myo-inositol and glycerol pools occurring later in the *X. elegans* drying response than they do in that of *X. schlechteri* and *X. humilis*.

This combination of both major physiological differences (poikilochlorophylly vs homoiochlorophylly) and subtle shifts in regulatory timing and sequencing within a clade whose overall stress response remains similar is an example of the plasticity and versatility of plant evolution.

### Proposing Pathway Regulation From Primary Metabolomes

While metabolite quantitation yields a “snapshot” of the biochemical state of a tissue, full understanding of physiological changes requires pathway flux information. In the absence of costly, and prohibitively complex isotope labelling experiments, changes in pathway flux may be estimated computationally. One such approach uses metabolomic data to infer changes in pathway activity in response to perturbations of the metabolic network such as the imposition of water deficit stress. Here, we have used the package PAPi to infer pathway regulation by mapping the most parsimonious combination of pathway up- and down-regulations that would result in the observed metabolite abundance changes (Aggio et al. 2010).

While this approach does not yield definitive answers, given many possible paths between the measured metabolic states, it can enrich the interpretation of metabolic data, and yield hypotheses for targeted transcriptomic or proteomic studies to test. In this case, the pathways listed in Table 2 are proposed as likely keys to the regulation of the desiccation response of the species surveyed.

Examination of the most highly regulated pathways reported in Table 2 shows some distinct patterns in relation to the desiccation tolerance strategies employed by the species surveyed. Among the RFO-dependent desiccation tolerant species, both starch and sucrose metabolism, and galactose metabolism were proposed to be among the most highly regulated pathways, reflecting the strong accumulation of both sucrose (from starch reserves) and galactosyl saccharides. By contrast, *M. flabellifolia* and *S. lepidophylla*, both trehalose-dependent resurrection plants, were proposed primarily to regulate galactose metabolism, reflecting their reduced dependency on sucrose accumulation.

The propanoate metabolism pathway was also strongly represented in regulation proposals for the trehalose-dependent species. This pathway is involved in the conversion of propanoic acid to succinyl- and acetyl-CoA and hence its direction to the TCA cycle. The presence of this pathway among those reported for the trehalose-dependent species may suggest a possible mechanism for the differential fitness of RFO-mediated desiccation tolerance and hence its wide adoption among flowering plants: Since propanoic acid is a product of lipid catabolism, it is plausible that RFOs may provide better protection for membranes and other lipid structures in the cell. This possibility is compatible with the proposed regulation of other lipid-related pathways in *M. flabellifolia* and *S. lepidophylla*, namely Butanoate metabolism and Fatty acid biosynthesis respectively.

Both *M. flabellifolia* and *C. pumilum* were proposed to regulate the metabolism of aromatic amino acids via the phenylalanine, tyrosine and tryptophan metabolic pathway. This provides a plausible mechanism for the accumulation of aromatic secondary metabolites, such as the polyphenol 3,4,5 tri- O-galloylquinic acid in *M. flabellifolia* (Moore et al. 2005).

The regulatory and metabolic difference between homoiochlorophyllous and poikilochlorophyllous resurrection plants is visible through the proposed differential regulation of the chlorophyll and porpohyrin pathway which includes the reaction for chlorophyll catabolism via the SGRL magnesium dechelatase (Shimoda et al. 2016).

Of particular interest are the proposed involvement of pathways previously little explored in the literature on desiccation tolerance. Several species were proposed to regulate the ABC transporter pathway. In plants, this suggests regulation of a wide variety of transporters, including sugar, amino acid and hormone transporters (Kretzschmar et al. 2011).

Additionally, a number of proposed pathway regulatory processes suggest – on the basis of the analysis of primary metabolomes alone – routes by which the accumulation of secondary metabolites associated with desiccation may be accumulated. These include the aforementioned aromatic amino acid pathways in *M. flabellifolia*, and the alkaloid, terpenoid and steroid pathways proposed for *Xerophyta* spp.

The prominence of inositol phosphate and related pathways in output for *M. flabellifolia* and *X.elegans* is interesting given recent research into the roles of inositol phosphate in abiotic stress signalling and the reconfiguration of plant lipids in response to stress (Munnik and Vermeer 2010; Munnik 2014), a topic that has received little direct attention in relation to resurrection plants.

### Data Variance And The Biophysical Constraints of Anhydrobiosis

Inspection of the PCA plots for each species suggests that the available state space for plants in anhydrobiosis is smaller than that in fully hydrated metabolism. That is to say, the small molecule complement of dry resurrection plants exist in a narrower, more defined band of concentrations than that in fully hydrated, unstressed plants. This observation was borne out by measures of the total variance of the entire hydrated and desiccated metabolome of each species (Fig. 14).

While the concept of statistical significance does not apply to measures of total variance, the observations are striking. Almost all angiosperm resurrection plants exhibited less variance in the dry state than in the hydrated. The desiccation-sensitive *A. thaliana* does not, and nor do the resurrection plants *S. lepidophylla* and *E. nindensis.* This observation is consistent with the hypothesis that the phase transition systems required for stable anhydrobiosis are constrained by the requirements of either vitrification (Buitink and Leprince 2004) or NADES formation (Dinakar and Bartels 2013), and that stable phase transitions (i.e. those that avoid crystallisation of micro- or macromolecules) can only occur within constrained portions of the chemical state space. In the cases of *S. lepidophylla* and *E. nindensis*, the constitutive nature of the DT strategy of the former is likely to mean that it always exists within an anhydrobiosis-competent state (Yobi et al. 2012), so convergence is not necessary, while *E. nindensis* comprises a mix of desiccation-sensitive and desiccation-tolerant tissues and throughout this study, exhibited very high levels of biological variation in every parameter measured. It is quite likely that sampling of presumed-tolerant tissues sometimes included some sensitive tissues and vice versa.

### Summary of Conclusions

Notwithstanding the chemical diversity within and between species, this study confirms that a limited number of core mechanisms are involved in enabling desiccation tolerance across the plant kingdom. All DT systems studied relied on either constitutively abundant trehalose, or the accumulation of raffinose family oligosaccharides and sucrose, with threshold ratios conditioned by other features of the metabolome. These core and highly conserved mechanisms suggest, if not a monophyletic origin for desiccation tolerance, that at least the necessary core mechanisms are very highly conserved, with trehalose biosynthesis being found across all biological kingdoms, and RFO biosynthesis being ancestral to most land plants.

The varied sequencing of different metabolic transitions, for instance the differences in timing of carbohydrate and amine shifts among *Xerophyta* spp, suggests that there is not a single “desiccation programme”, but that different subprocesses are coordinated at different phases of drying, as mild water deficit progresses to severe water loss and desiccation. This is likely highly influenced by the nature of shut down of photosynthesis, i.e. homoiochlorophylly vs poikilochlorophylly, with the former continuing this metabolism to lower RWC’s than the latter (Farrant and Hilhorst 2022).

Achieving anhydrobiosis and longevity in the dry state involves the simultaneous regulation of many metabolic processes, and the convergence on a limited range of desiccation-competent compositions. This study has shown that there are likely to be constraints on the composition of a viable dry state, and has yielded some insights into how different adaptive strategies interact with the biophysical constraints of anhydrobiosis, however a full understanding of the cellular biophysics involved will require a more detailed understanding of the phase systems involved, including how the small molecule complement interacts with the macromolecular environment.

## Author contributions

The study was conceptualised by HJWD, JMF and HWMH.

JMF facilitated collection of the resurrection plants used in this study, gave intellectual advice and contributed towards editing of the article.

## Acknowledgments

JMF acknowledges funding from the Department of Science and Innovation, National Research Foundation South African Research Chair grant number 98406.

## Data availability statement

### Supporting information

Additional supporting information may be found online in the Supporting Information section at the end of the article**. Table S1, Table S2, Figure S1, Figure S2, Figure S3, Figure S4.**

