## Supplemental tables and figures for "A Horizontal View of Primary Metabolomes in Vegetative Desiccation Tolerance"

### Supplementary Data

\#

Table S1: Abundance changed and p-values of quantitated sugars in each species:

|  |  | **Wilted:Hydrated** | | **Dry:Wilted** | | **Dry:Hydrated** | |
| --- | --- | --- | --- | --- | --- | --- | --- |
| **Species** | **Sugar** | **Fold-change** | **p-value** | **Fold-change** | **p-value** | **Fold-change** | **p-value** |
| *Arabidopsis thaliana* | Glucose |  |  |  |  | 0.31 | 0.1254 |
|  | Fructose |  |  |  |  | 1.80 | 0.56956 |
|  | Sucrose |  |  |  |  | 2.88 | 0.0001 |
|  | Trehalose |  |  |  |  | 1.22 | 0.4941 |
|  | Raffinose |  |  |  |  | 0.2 | 0.0890 |
|  | Stachyose |  |  |  |  | n.d. | n.d. |
| *Craterostigma pumilum* | Glucose | 2.55 | 0.3773 | 0.10 | 0.1588 | 0.24 | 0.7693 |
|  | Fructose | 1.53 | 0.8833 | 1.48 | 0.7897 | 2.26 | 0.5229 |
|  | Sucrose | 21.67 | 0.1092 | 2.77 | 0.0092 | 60 | 0.0010 |
|  | Trehalose | 1.17 | 0.8894 | 0.70 | 0.6311 | 0.82 | 0.8806 |
|  | Raffinose | 1.27 | 0.6728 | 1.71 | 0.8376 | 2.18 | 0.3752 |
|  | Stachyose | 2.69 | 0.5809 | 0.71 | 0.8849 | 1.92 | 0.8431 |
| *Eragrostis nindensis (Non-Senescent)* | Glucose | 2.17 | 0.3019 | 0.32 | 0.1047 | 0.69 | 0.8721 |
|  | Fructose | 3.92 | 0.2398 | 0.26 | 0.1352 | 1.03 | 0.9997 |
|  | Sucrose | 10.41 | 0.0796 | 0.69 | 0.3680 | 7.17 | 0.1806 |
|  | Trehalose | 2.9 | 0.2391 | 0.23 | 0.0922 | 0.68 | 0.9269 |
|  | Raffinose | 107.5 | 0.6999 | 1.29 | 0.9400 | 139 | 0.5410 |
|  | Stachyose | (inf) | 0.7452 | 0.76 | 0.9742 | (inf) | 0.8096 |
| *Eragrostis nindensis (Senescent)* |  |  |  |  |  |  |  |
|  | Glucose | 3.14 | 0.0273 | 0.77 | 0.1486 | 2.44 | 0.0397 |
|  | Fructose | 3.12 | 0.1286 | 0.66 | 0.2837 | 3.26 | 0.2809 |
|  | Sucrose | 1.85 | 0.9844 | 1.82 | 0.9304 | 3.38 | 0.8455 |
|  | Trehalose | 3.68 | 0.8652 | 0.79 | 0.9804 | 2.90 | 0.8951 |
|  | Raffinose | 4.66 | 0.1750 | 0.41 | 0.2004 | 1.91 | 0.6943 |
|  | Stachyose | n.d. |  | n.d. |  | n.d. |  |
| *Eragrostis tef* | Glucose | 8.41 | 0.0125 | 0.36 | 0.0472 | 3.0 | 0.5187 |
|  | Fructose | 8.13 | 0.0065 | 0.31 | 0.0199 | 2.53 | 0.5742 |
|  | Sucrose | 1.77 | 0.0216 | 1.79 | 0.0011 | 3.16 | 0.0001 |
|  | Trehalose | 0.94 | 0.9677 | 1.32 | 0.4723 | 1.24 | 0.6039 |
|  | Raffinose | 3.26 | 0.1850 | 1.25 | 0.7543 | 4.08 | 0.0729 |
|  | Stachyose | (inf) | 0.1821 | 1.07 | 0.9899 | (inf) | 0.1533 |
| *Myrothamnus flabellifolia* | Glucose | 0.29 | 0.0079 | 0.50 | 0.5116 | 0.14 | 0.0019 |
|  | Fructose | 1.11 | 0.9787 | 0.36 | 0.4511 | 0.40 | 0.4835 |
|  | Sucrose | 51.56 | 0.0159 | 1.07 | 0.9469 | 55.16 | 0.0074 |
|  | Trehalose | 0.79 | 0.6325 | 1.10 | 0.9298 | 0.87 | 0.7956 |
|  | Raffinose | 13.6 | 0.0306 | 0.61 | 0.3341 | 8.24 | 0.1307 |
|  | Stachyose | 34.5 | 0.0245 | 0.22 | 0.0579 | 7.65 | 0.6749 |
| *Selaginella lepidophylla* | Glucose |  |  |  |  | 0.026 | 1.25e-5 |
|  | Fructose |  |  |  |  | 1.68 | 0.6939 |
|  | Sucrose |  |  |  |  | 35.45 | 0.0149 |
|  | Trehalose |  |  |  |  | 0.81 | 0.4600 |
|  | Raffinose |  |  |  |  | (inf) | 0.2963 |
|  | Stachyose |  |  |  |  | (inf) | 0.03 |
| *Xerophyta elegans* | Glucose | 1.0 | 1.0 | 0.055 | 0.0441 | 0.055 | 0.0438 |
|  | Fructose | 20 | 0.0053 | (-inf) | 0.0002 | (-inf) | 0.0326 |
|  | Sucrose | 4.20 | 0.0280 | 2.40 | 0.0000 | 10.07 | 0.0008 |
|  | Trehalose | 0.17 | 0.9087 | 13.1 | 0.5508 | 2.18 | 0.8043 |
|  | Raffinose | 8.19 | 0.2986 | 1.36 | 0.7688 | 11.01 | 0.1003 |
|  | Stachyose | (inf) | 0.1903 | 2.15 | 0.1232 | (inf) | 0.0078 |
| *Xerophyta humilis* | Glucose | 0.17 | 0.0207 | 0.40 | 0.8836 | 0.070 | 0.0122 |
|  | Fructose | 0.28 | 0.0175 | 0.64 | 0.8456 | 0.18 | 0.0095 |
|  | Sucrose | 3.63 | 0.0033 | 1.04 | 0.9603 | 3.76 | 0.0026 |
|  | Trehalose | 0.49 | 0.4541 | 1.68 | 0.6946 | 0.82 | 0.8992 |
|  | Raffinose | 2.80 | 0.0136 | 1.33 | 0.1589 | 3.72 | 0.0017 |
|  | Stachyose | 6086 | 0.0198 | 0.99 | 0.9997 | 6047 | 0.0203 |
| *Xerophyta schlechteri* | Glucose | 0.068 | 3.6e-6 | 0.50 | 0.7874 | 0.034 | 3.0e-6 |
|  | Fructose | 0.33 | 0.0045 | 0.69 | 0.7133 | 0.22 | 0.0022 |
|  | Sucrose | 3.94 | 0.0056 | 0.84 | 0.5702 | 3.32 | 0.0168 |
|  | Trehalose | n.d. |  | n.d. |  | n.d. |  |
|  | Raffinose | 11.89 | 0.0039 | 0.77 | 0.4172 | 9.19 | 0.01457 |
|  | Stachyose | 84.5 | 0.0234 | 1.0 | 1.0 | 84.5 | 0.0233 |

Table S2: Annotated GCMS peaks

\#

| **Retention Time / min** | **Base Peak m/z** | **Annotation** |
| --- | --- | --- |
| 15.5162857 | 174 | 4-aminobutanoic acid (GABA) |
| 8.25114286 | 57 | alkane 10 |
| 12.72625 | 57 | alkane 13 |
| 14.00825 | 57 | alkane 14 |
| 15.2224286 | 57 | alkane 15 |
| 16.3642 | 57 | alkane 16 |
| 20.4181667 | 57 | alkane 20 |
| 21.2491429 | 57 | alkane 21 |
| 22.7183333 | 57 | alkane 23 |
| 23.446 | 57 | alkane 24 |
| 24.17575 | 57 | alkane 25 |
| 26.994 | 57 | alkane 29 |
| 11.3367 | 228 | artefact |
| 16.3755 | 188 | asparagine |
| 17.433 | 188 | asparagine 2 |
| 17.8 | 205 | aspartic acid |
| 18.4778333 | 273 | citric acid |
| 7.968 | 157 | cyclohexanol |
| 21.0425 | 174 | dopamine |
| 15.625 | 292 | erythronic acid |
| 12.1394286 | 147 | ethylmalonic acid |
| 18.923 | 103 | fructose |
| 19.023375 | 217 | fructose 2 |
| 27.3475 | 204 | galactinol |
| 19.872 | 458 | gallic acid |
| 26.348 | 361 | gentiobiose |
| 22.39 | 333 | glucoheptonic acid |
| 17.94625 | 217 | glucopyranose -H2O |
| 19.938 | 204 | Glucopyranose, D- (5TMS) |
| 19.2925 | 204 | Glucopyranoside, 1-O-methyl-, beta-D- (4TMS) |
| 19.198125 | 319 | glucose |
| 19.408625 | 147 | glucose 2 |
| 12.9255 | 189 | glyceric acid |
| 12.161 | 205 | glycerol |
| 12.7376 | 174 | glycine |
| 10.0238 | 133 | hydroxylamine |
| 18.57 | 147 | isocitric acid |
| 12.533 | 158 | isoleucine |
| 9.0682 | 117 | lactic acid |
| 14.91325 | 147 | malic acid |
| 21.02825 | 217 | myo-inositol |
| 22.242 | 117 | octadecanoic acid |
| 16.761 | 218 | phenylalanine |
| 17.65 | 293 | phenylpyruvic acid |
| 12.214625 | 299 | phosphoric acid |
| 15.5444 | 156 | pyroglutamic acid |
| 18.8223333 | 345 | quinic acid |
| 28.5178 | 345 | Quinic acid, 3-caffeoyl-, E- |
| 31.05125 | 361 | raffinose |
| 17.438375 | 217 | ribitol |
| 21.148 | 315 | ribose-5-phosphate |
| 12.115 | 114 | serine |
| 22.2307143 | 147 | succinic acid |
| 24.837625 | 361 | sucrose |
| 12.612 | 218 | threonine |
| 25.6126667 | 361 | trehalose |
| 22.54 | 204 | unknown disaccharide |
| 11.4026667 | 144 | valine |
| 12.65 | 142 | proline |

\#

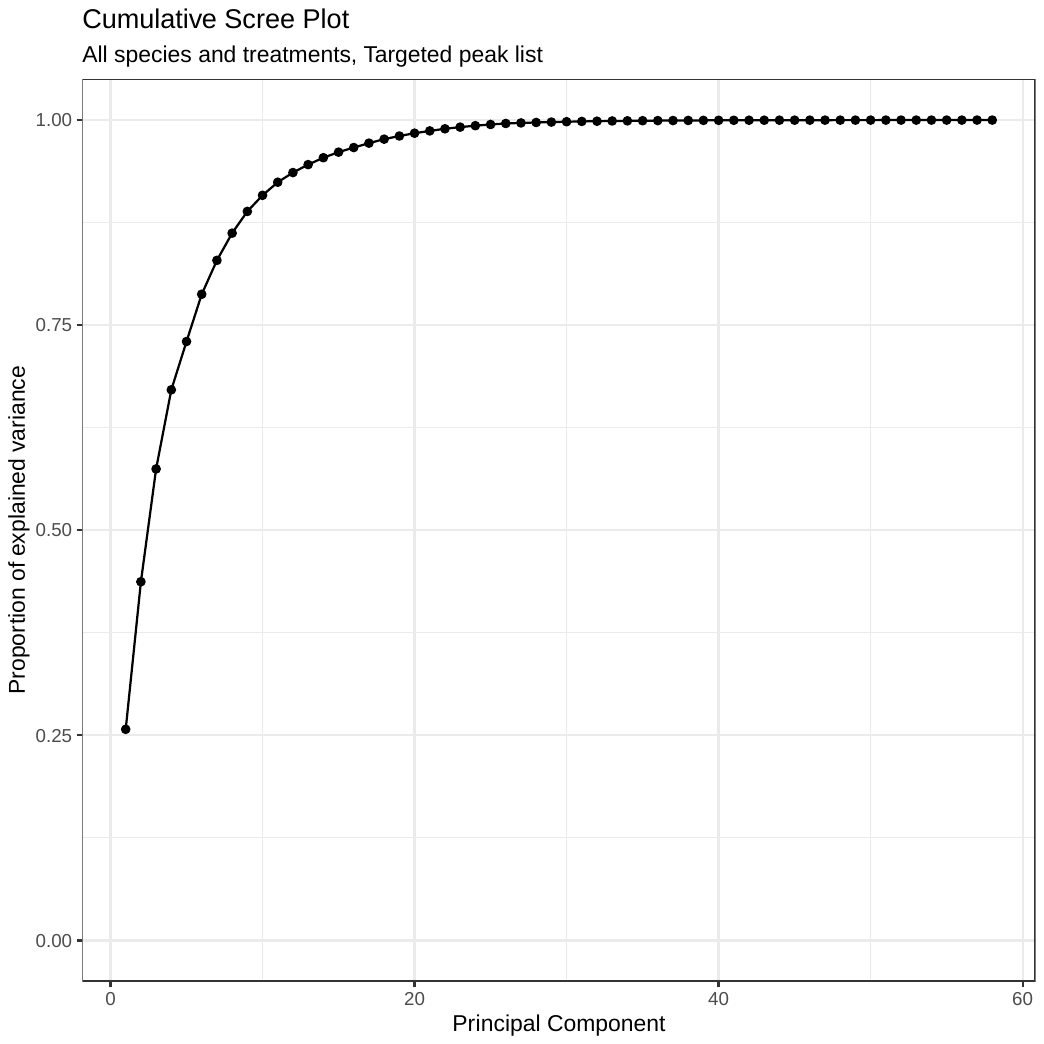

Figure S1

Screeplot of variance per principal component for PCA analysis of targeted GCMS peak list.

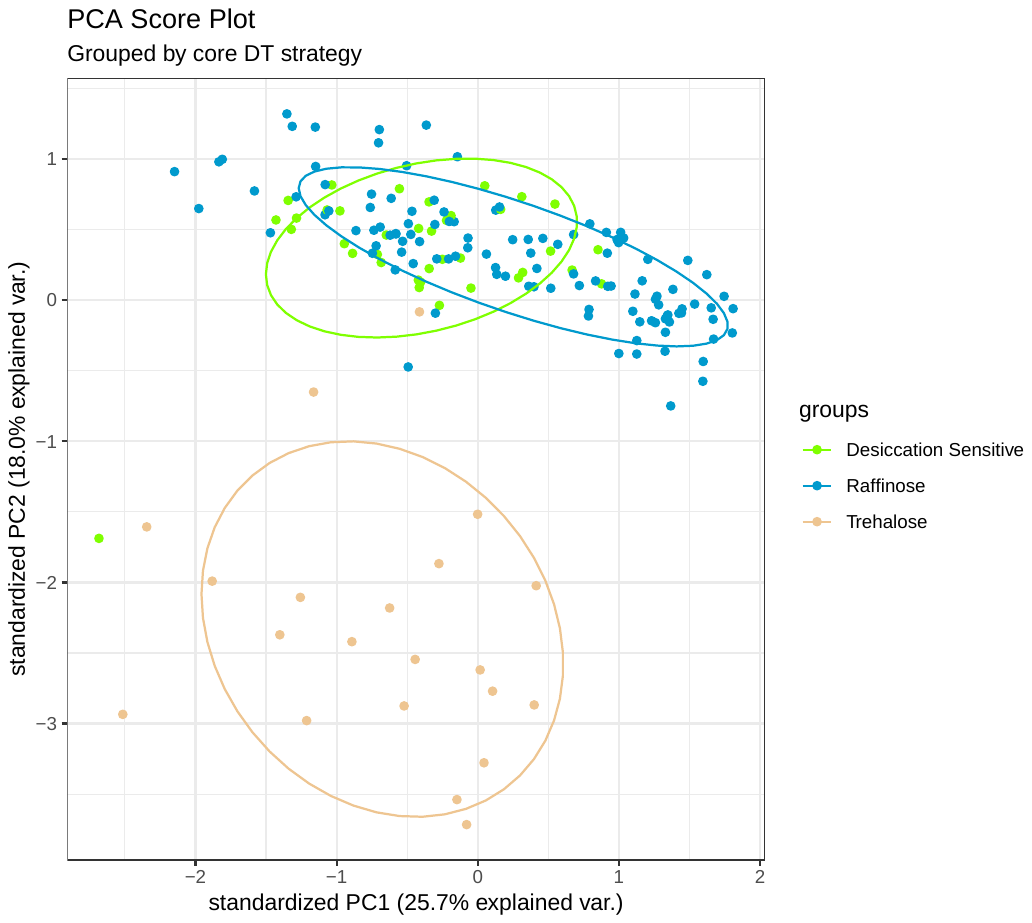

Figure S2

Score plot of principal components 1 and 2 for PCA analysis of targeted GCMS peak list, coloured by core DT strategy.

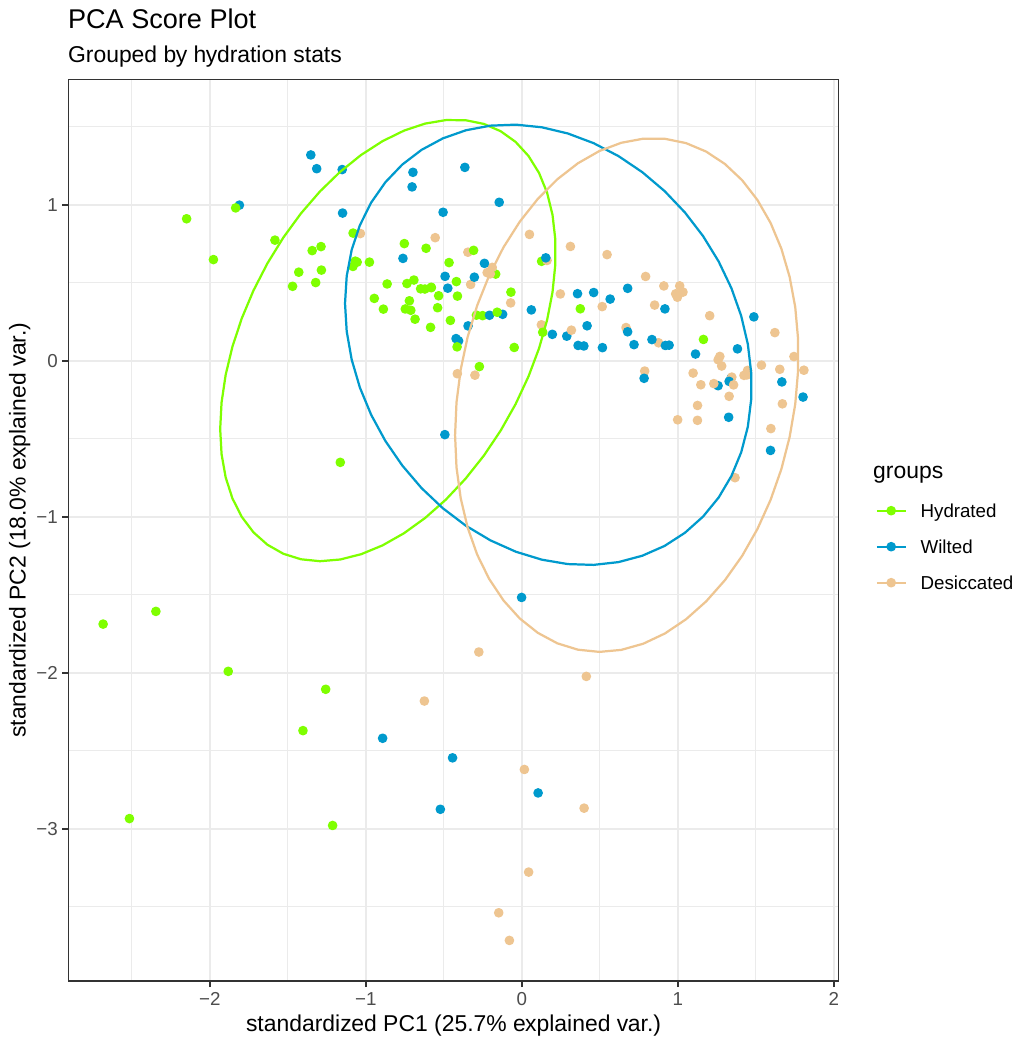

Figure S3

Score plot of principal components 1 and 2 for PCA analysis of targeted GCMS peak list, coloured by sample hydration status.

Figure S4

Per-species PCA analysis of targeted GCMS peak list.
